# Improved circulating tumor DNA profiling by simultaneous extraction of DNA methylation and copy number information from Methylated DNA Sequencing data (MeD-seq)

**DOI:** 10.1101/2025.01.21.633371

**Authors:** Daan M Hazelaar, Ruben G Boers, Joachim B Boers, Vanja de Weerd, Jean Helmijr, Maurice PHM Jansen, Henk MW Verheul, Cornelis Verhoef, Joost Gribnau, John WM Martens, Stavros Makrodimtris, Saskia M Wilting

## Abstract

Cell-free DNA (cfDNA) analysis offers a powerful, minimally invasive approach to improve cancer care by measuring tumor-specific genomic and epigenetic alterations. Here, we demonstrate the versatility of MeD-seq, a methylation-dependent sequencing assay, for comprehensive cfDNA analysis, including DNA methylation profiling, chromosomal copy number (CN) alterations, and tumor fraction (TF) estimation. MeD-seq-derived CN profiles and TF estimates from 38 colorectal cancer with liver metastases (CRLM) and 5 ovarian cancer patients were highly comparable to shallow whole-genome sequencing (sWGS) validating our approach. For 120 CRLM patients we used MeD-seq CN and TF information in an improved Differential Methylation Model which detected additional significantly Differentially Methylated Regions (DMRs) correlating with TF estimates. Using the identified DMR sets we were subsequently able to distinguish healthy blood donors from CRLM patients with low amounts of circulating tumor DNA (ctDNA) as well. These findings show MeD-seq as an affordable platform for detecting cancer-specific signals directly from plasma without prior tissue-based information. Future work could expand its application to other cancer types, solidifying MeD-seq as a versatile tool for cfDNA profiling.

**Graphical abstract:** Graphical Abstract of MeD-seq and Copy Number Informed Differential Methylation Analysis
Liquid biopsy samples are collected, followed by cfDNA extraction, methylation-dependent enzymatic digestion, Whole Genome Sequencing (WGS) and *in silico* read selection. Background reads (lacking the recognition site of the methylation-dependent restriction ezyme) are utilized for CN profiling and TF estimation. Methylated reads (with the recognition site of the methylation-dependent restriction enzyme) are counted in CpG islands and integrated with CN information to perform Differential Methylation profiling, enabling a comprehensive assessment of methylation in cfDNA.

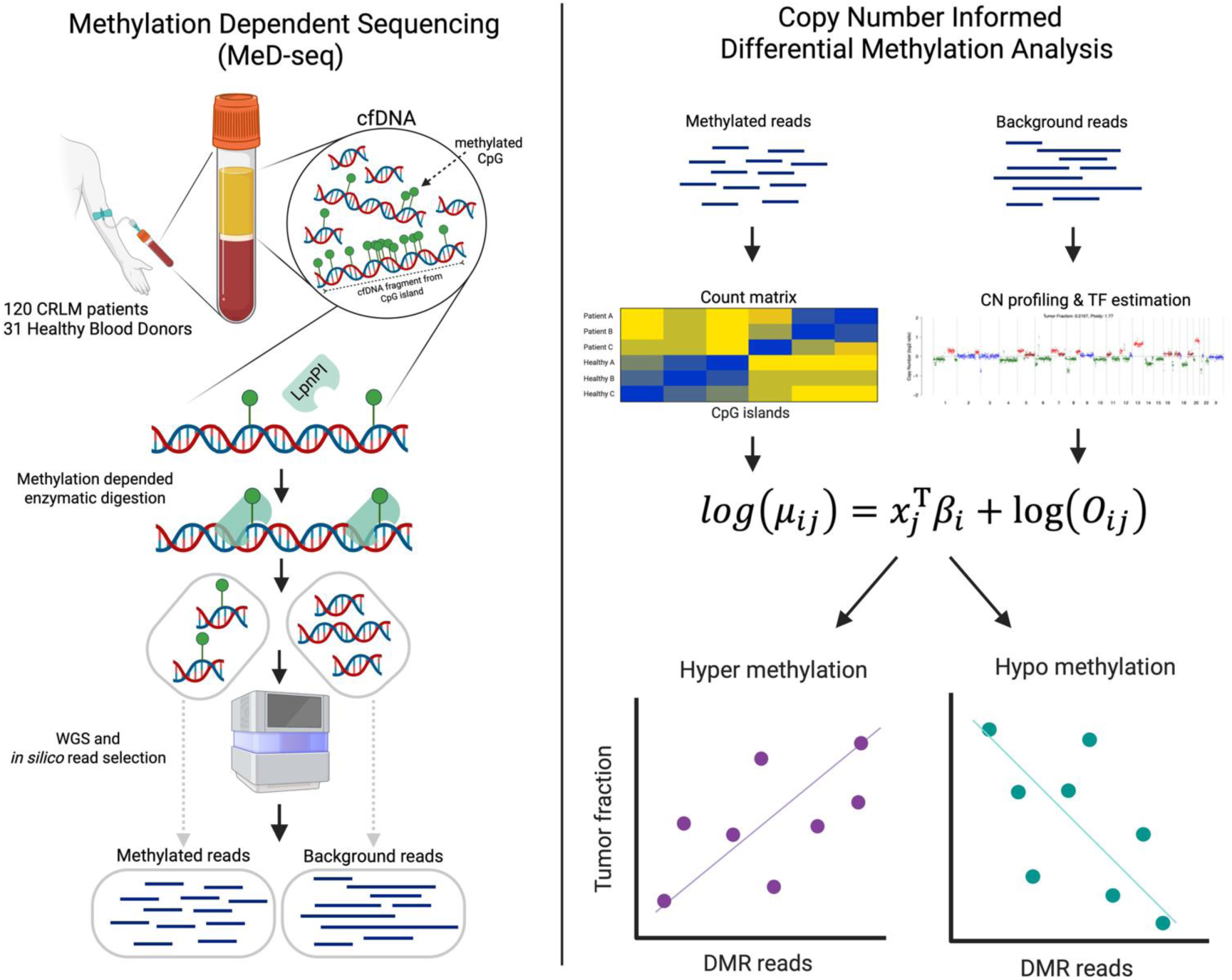

## Introduction

Over the past decade liquid biopsies, i.e. the utilization of bodily fluids to detect disease-specific signals, has surged in cancer research. Specifically, circulating cell-free DNA (cfDNA) fragments in the blood stream originating from succumbed cells provide a promising means to obtain crucial information on their cells of origin (Wan et al., 2017). While not yet widely integrated into standard diagnostic or disease monitoring protocols, liquid biopsies hold significant potential for early detection and tracking of disease progression, prediction of prognosis, and treatment efficacy (Connal et al., 2023; Penny et al., 2024; Reichert et al., 2023).

To distinguish between circulating tumor DNA (ctDNA) and cfDNA from healthy cells, one can focus on a number of general oncogenic hallmarks present in ctDNA such as tumor-specific mutations (André et al., 2024), chromosomal alterations (Adalsteinsson et al., 2017), altered DNA methylation profiles (Y. Liu et al., 2024; Zhou et al., 2022), and fragmentomics features (Mathios et al., 2021; Moldovan et al., 2024; Mouliere et al., 2018). Previous studies have highlighted the potential of cfDNA analysis in cancer research, demonstrating its utility for clinical predictions, cancer subtyping, and identifying biological signatures across diverse cancer types (Cristiano et al., 2019; Doebley et al., 2022). However, the limited amount of cfDNA in a blood sample restricts the range and number of assays one can perform on a single sample. Moreover, these assays are often costly, making it unfeasible to perform all of them even if sufficient material were available.

To overcome these limitations, bioinformatics efforts are being developed to allow multimodal read-outs from individual assays with the goal to conserve patient material, reduce costs, and improve ctDNA detection (Katsman et al., 2022; J. Liu et al., 2024; Y. Liu et al., 2024; Nguyen et al., 2023; Thierry, 2023; Wang et al., 2023). Inspired by these initiatives, we here investigate whether we can extract additional, relevant information from the data generated by our previously described Methylated DNA Sequencing (MeD-seq) assay for cfDNA methylation profiling (Deger et al., 2021).

MeD-seq uses a methylation-dependent restriction enzyme (LpnPI) to digest methylated (cf)DNA into 32bp fragments, which are amendable after size selection for next generation sequencing (Graphical abstract). The sequenced fragments that contain an LpnPI recognition site at the expected position —methylated reads— are then mapped to the human genome to associate each of them with a CpG site after which a methylation profile can be constructed (Boers et al., 2023) (Graphical abstract). This approach offers several key advantages over other cfDNA methylation profiling methods: a) the broad coverage of CpG sites (>50% genome wide), b) the lack of need for chemical conversion of methylated DNA, c) its low DNA input requirement (5-10ng), d) its low costs (only 20-30M reads needed to all CpGs), and e) the short hands-on time (Boers et al., 2018; Deger et al., 2021).

The original MeD-seq computational pipeline focuses on methylated reads and ignores reads that do not contain an LpnPI recognition site —background reads— as these are not directly informative for determining the methylation status of the DNA (Boers et al., 2018). In practice, this means only a median of 37% of all sequencing reads derived from cfDNA using the current MeD-seq protocol are used (Deger et al., 2021). We hypothesized that these ignored background reads represent the sample’s cfDNA fragment distribution prior to digestion and may thus provide information on the chromosomal coverage of the sample which can be used to extract additional information as well as improve differential methylation analysis of cfDNA.

Here, we aim to utilize background reads from MeD-seq data to construct sample-specific chromosomal Copy Number (CN) profiles and estimate Tumor Fraction (TF). Then, we incorporate this information into a Differential Methylation Model to improve the detection of cancer-specific Differentially Methylated Regions (DMRs) in cfDNA.

## Materials and Methods

### Patient and control samples

We used blood samples from five consenting patients with metastatic ovarian cancer and 120 consenting patients with colorectal liver metastases (CRLM) (43 females and 70 males, age: 38-84). From the CRLM patients 139 blood samples were evaluated, 120 were collected preoperatively (T0) and 19 postoperatively (T3). The clinical studies from which the used samples were obtained were both approved by the Medical Ethics Committee of the Erasmus University Medical Center (MEC 15-289, and MEC 17-238). In addition, 10 ml of blood was obtained from 51 consenting anonymous healthy blood donors (HBDs) (22 females and 29 males, age: 22-71) via the Dutch National blood bank (Sanquin).

All blood samples were collected in CellSave tubes and plasma was isolated from the obtained blood within 96 h after blood draw by 2 sequential centrifugation steps at room temperature (10 min at 1711 g followed by 10 min at 12,000 g) and stored at − 80 °C (van Dessel et al., 2017). cfDNA from plasma was extracted using the QiaAmp kit (Qiagen) according to manufacturer’s instructions. cfDNA concentration was determined using the Quanti-IT dsDNA high-sensitivity Assay (Invitrogen) and stored at -30 °C until further handling.

We performed sWGS and MeD-seq on cfDNA isolated from the same blood sample for five ovarian cancer patients and 38 samples from 19 CRLM patients (6 female, 13 male) at T0 and T3. For the remaining 101 CRLM patients, MeD-seq analysis was only conducted on the T0 sample. For the HBDs, sWGS was performed on cfDNA from 20 individuals (10 female, 10 male), while MeD-seq analysis was performed on a different set of 31 healthy individuals (12 female, 19 male).

### MeD-seq and sWGS analyses of cfDNA

MeD-Seq was performed as previously described (Boers et al., 2018; Deger et al., 2021). In short, 10ng of cfDNA was digested with LpnPI (New England Biolabs, Ipswich, MA) yielding 32 bp fragments around the fully methylated CpG containing recognition site (motif: CCGG, GCGG, CCGA, ACGG, CCGC, CCGT, TCGG, GCGC). Samples were prepared for sequencing using the ThruPLEX DNA-seq 96D kit and the ThruPLEX DNA-Seq HV kit (Rubicon Genomics, Takara Bio Europe, Saint-Germain-en-Laye, France), for CRLM and ovarian cancer samples respectively, and purified on a Pippin HT system with 3% agarose gel cassettes (Sage Science, Beverly, MA) enriching for fragments ranging from 148 to 192bp (including sequencing adapters), which was shown to enrich for tumor-derived DNA fragments (Mouliere et al., 2018). Libraries were multiplexed and sequenced until a total of ∼20M reads on either an Illumina HiSeq 2500 for 50 bp single-end reads (CRLM samples) or an Illumina NEXTseq2000 (ovarian cancer) for 50 bp paired-end reads, according to the manufacturer’s instructions (Illumina, San Diego, CA).

For sWGS, indexed sequencing libraries were prepared using the ThruPLEX DNA-Seq HV kit with an input of 5 ng of cfDNA per sample. Libraries were sequenced to a median 1.0x depth of coverage on an Illumina NEXTseq2000, generating at least 10M 150 bp paired-end reads. For ovarian cancer the combined shWGS and exome-seq workflow from Twist BioScience was applied, although the exome data was not used in this study.

### Quality control, pre-processing and read alignment

We used Fastp (v0.23.4) for adapter removal and deduplication, and BWA-MEM (v0.7.18) with soft-clipping enabled to align reads to the hg38 reference genome. Next, duplicate reads were removed using Sambamba (v1.0.1) markdup, as well as reads that mapped to blacklisted regions (ENCODE v2) and reads with mapping quality below 5.

After preprocessing, MeD-seq reads were separated based on the presence of an LpnPI recognition site located 12–16 bp from the start of the read. Reads containing the LpnPI recognition site are hereafter referred to as methylated reads, while those lacking this signal are referred to as background reads.

### Construction of CNV profiles

To evaluate chromosomal alterations, we employed ichorCNA, an established CNV calling method for sWGS data of cfDNA (Adalsteinsson et al., 2017). For sWGS data, cfDNA fragments up to 150 bp were selected in silico, which was shown to enrich for tumor-derived DNA fragments (Mouliere et al., 2018). For copy number analysis using MeD-seq data, we used the background reads as input for ichorCNA.

For both assays, the genome was binned into intervals of 1Mbp, and the number of reads in each bin was counted. Bins with a copy number gain or loss are expected to have more or fewer reads, respectively, compared to a diploid genome. ichorCNA employs a set of cfDNA profiles from healthy individuals (panel of normals) to estimate the expected read counts in each bin. To avoid potential biases due to the use of a different assay, we constructed assay-specific panels of normals using either MeD-seq background reads (n=31) or sWGS data (n=20). Within each sample, each bin’s read count was corrected for GC content and mappability biases as described in (Adalsteinsson et al., 2017). Subsequently, the log ratio of the corrected read count in each bin to the corresponding median corrected read count in the appropriate panel of normals was calculated. These log ratios were then used as input for the ichorCNA Hidden Markov Model, which returns the most likely copy number state of each bin as well as an estimate of the tumor fraction (Adalsteinsson et al., 2017).

We used default parameters for the ichorCNA model (v0.3.2) except that chromosome 19 was excluded from the training phase as recommended by the authors, and the model was initialized using two prior normal fractions of 0.5 and 0.6 and the result with the highest likelihood was selected. Additionally, for MeD-seq samples, we adapted the ichorCNA source code to include sex as a parameter as determining the sex from the binned data showed to be inconsistent.

### Evaluation of TF estimates and CN-profiles

To assess systematic bias in the ichorCNA data and model we employed Bland-Altman (BA) analysis (Altman & Bland, 1983; Giavarina, 2015). In short, Bland-Altman analysis is typically used to compare two continuous measurements of the same quantity by plotting their average on the x-axis and the difference between them on the y-axis. Additionally, we calculated summary statistics, including the mean difference between the measurements and the 95% confidence intervals for both the data points and the mean difference, using bootstrapping (n=10,000). Additionally, we used Spearman correlation to compare TF estimates and ichorCNA logR ratios based on MeD-seq and sWGS data.

### Agreement between CN-profiles

The ichorCNA model assigns a discrete state to each bin (ranging from 1-5; with 1 being hemizygous loss and 5 high-level amplification), representing the copy number state of that bin. We quantified the agreement between the copy number states estimated from MeD-seq and sWGS data from the same sample using Cohen’s Kappa.

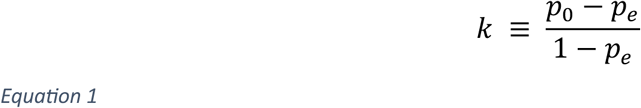

With *k* as the value for kappa, bounded between -1 (perfect disagreement) and 1 (perfect agreement), *p*_0_ as the observed agreement and *p_e_* as the probability of a by chance agreement.

### Association between noise and observed agreement in CN-profiles

A low sequencing depth leads to noisy coverage profiles which can have a detrimental effect on the detection of CNVs. To quantify noise within a CN-profile, we calculated the Median Absolute Deviation (MAD) of each CN-profile by:

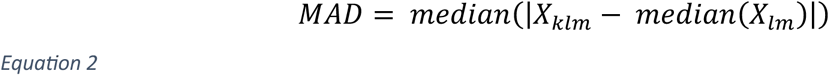

With, X as the log ratio of bin *k* in CN segment *l* in sample *m*.

To determine if differences in noise levels between MeD-Seq and WGS-derived CN profiles could be explained by differences in coverage, we down-sampled the BAM files of five high-coverage ovarian cancer sWGS samples to 75%, 50%, 25%, 10%, and 5% of the original read count and compared the MAD scores of these down-sampled samples and their MeD-seq counterparts.

### Differential methylation analysis

For differential methylation analysis we counted the methylated reads in 31 HBDs and 120 CRLM patients, not considering the background reads, mapping to each of the 27,923 CpG islands (Perez et al., 2024). Islands with low counts across most samples are prone to false positive differential methylation calls as a few outliers can have large influence on the mean, so we removed CpG islands with fewer than 10 counts per million in more than 70% of the samples in both groups. Additionally, we determined the coefficient of variation for each CpG island keeping only the 50% most variable features in chromosomes 1-22 and X.

MeD-seq derived counts exhibit overdispersion, meaning extra variability that exceeds what is expected under a simple Poisson model, due to both biological and technical variability. This characteristic of the data makes the negative binomial distribution a suitable choice for our Differential Methylation Model. We used negative binomial models implemented in the edgeR package (version 4.2.2), which has also been used for processing cfDNA methylation data generated using the MEDIP-seq assay (Lienhard et al., 2014). More specifically, we modelled the observed count *y_ij_* of reads mapped to a specific CpG island *i* in sample *j*. The model is defined as:

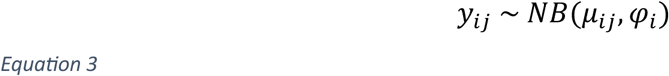

With *μ_ij_* as the expected count for CpG island *i* and sample *j* given the total number of methylated reads in sample *j,* and *φ_i_* as the dispersion parameter for CpG island *i.* The dispersions were estimated by maximizing the negative binomial likelihood using weighted likelihood empirical Bayes to obtain posterior dispersion estimates as is standard in edgeR analysis (Phipson et al., 2016). Assuming an estimate for *φ_i_* is available, the logarithm of mean *μ_ij_* can be fitted using a log linear model:

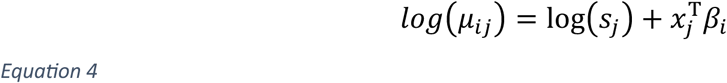

With *s_j_* as the library size normalization factor estimated using the TMM method (Robinson & Oshlack, 2010), and *x_j_* the design matrix containing covariates: *age*, *sex* and an indicator variable denoting whether a sample is from a cancer patient or a healthy control. The vector

of regression coefficients for each covariate associated with CpG island *i* is denoted by *β_i_*. After model fitting, we used edgeR’s Quasi-Likelihood F-Test to evaluate differential methylation between HBDs and CRLM samples (Chen et al., 2016). This tests the null hypothesis that the β coefficient corresponding to the sample type (healthy or cancer) is equal to 0 resulting in a p-value (FDR-adjusted, Benjamini-Hochberg) and corresponding log Fold Change (logFC) for each CpG island.

To correct for changes in the expected counts caused by local copy number changes, we adapted the model above by adding an offset matrix *O_ij_*:

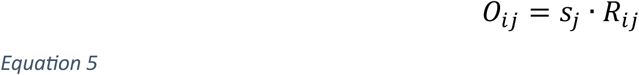

where *R_ij_* is the ratio of observed background reads in sample j to the median of the healthy samples in the genomic bin containing CpG island *i*, as determined using ichorCNA. We imputed missing ratios using Kalman smoothing (imputeTS, v3.3) for gaps of up to 5 bins. We then replaced the *s_j_* term in equation 4 with *O_ij_* and repeated the model fitting and hypothesis tests as described before.

To reduce the influence of outlier read counts on DMR detection, we bootstrapped the entire analysis procedure 1,000 times. We summarized the logFC and p-values across all bootstrap iterations by taking their median values. The resulting median p-values were FDR-adjusted using the Benjamini–Hochberg method, and CpG islands with an adjusted p-value <0.05 were considered a significantly Differentially Methylated Region (DMR).

We repeated this analysis using a) the entire set of 120 CRLM patients, b) only patients with high ctDNA load based on the CNV profiles and c) only those with low ctDNA load. To ensure fair comparisons between the different sample subsets, we matched the number of cancer samples used for model fitting in each bootstrap iteration, thereby maintaining equal statistical power across the subsets.

### DMR validation

To internally validate the tumor-specificity of the identified DMRs, we calculated the Spearman correlation between tumor fraction (TF) estimates and the normalized, log-transformed counts per million (CPM) for each CpG island, using only the high-TF CRLM samples. For CpG islands labeled as DMRs, we performed a one-sided test based on the direction of the corresponding logFC: testing for a positive correlation in hypermethylated regions and a negative correlation in hypomethylated regions. For CpG islands not identified as DMRs, a two-sided test was applied. All resulting p-values were FDR-corrected using the Benjamini–Hochberg method, and correlations with adjusted p-values <0.05 were considered statistically significant. Additionally, we determined the fraction of methylated reads that were mapped to a DMR as an indicator for tumor load and compared this with IchorCNA TF estimates.

### Implementation details

We developed a freely available data processing pipeline, called nf-core-medseqcn, that starts from the fastq files from sWGS or Med-seq. To ensure our pipeline is portable, scalable, and can be applied reproducibly we used the nf-core template of Nextflow (Di Tommaso et al., 2017; Ewels et al., 2020). All components of the pipeline are provided as Docker and Singularity containers through the Biocontainer’s initiative (da Veiga Leprevost et al., 2017). The pipeline includes multiple components for quality control, alignment, processing, and fitting the ichorCNA model for both MeD-seq and sWGS sequencing data. It produces a structured output to support subsequent downstream analyses and is available at https://github.com/daanhazelaar/nf-core-medseqcn.

### Data availability

All processed data—including copy number profiles, methylated read counts, DMR profiles, reference files, and R objects required to reproduce the results and figures—are available on on zenodo: <u>10.5281/zenodo.15438260</u>.. All analysis scripts are available at https://github.com/daanhazelaar/med-seq-ctDNA-profiling/

## Results

### Enrichment of MeD-seq background around LpnPI recognition sites alleviated by bias correction

To assess whether MeD-seq background reads contain information on chromosomal alterations, we evaluated sequencing metrics from 38 CRLM and five ovarian cancer cfDNA samples to confirm sufficient coverage (mean: 10.5%, min: 2.5%, max: 19.2%, Figure 1A) and read mapping quality (mean: 42.5, min: 37.8, max: 53.1) (Figure 1B, Supplemental Figure 1). Next, we analyzed the impact of LpnPI digestion on the genomic distribution of methylated and background reads, revealing increased LpnPI recognition site frequencies at +12 bp and - 12 bp from the start and end of sequencing reads for both read types (Supplemental Figure 2). This pattern, absent in sWGS reads, suggests that MeD-seq background reads are partially influenced by LpnPI digestion.

**Figure 1.**
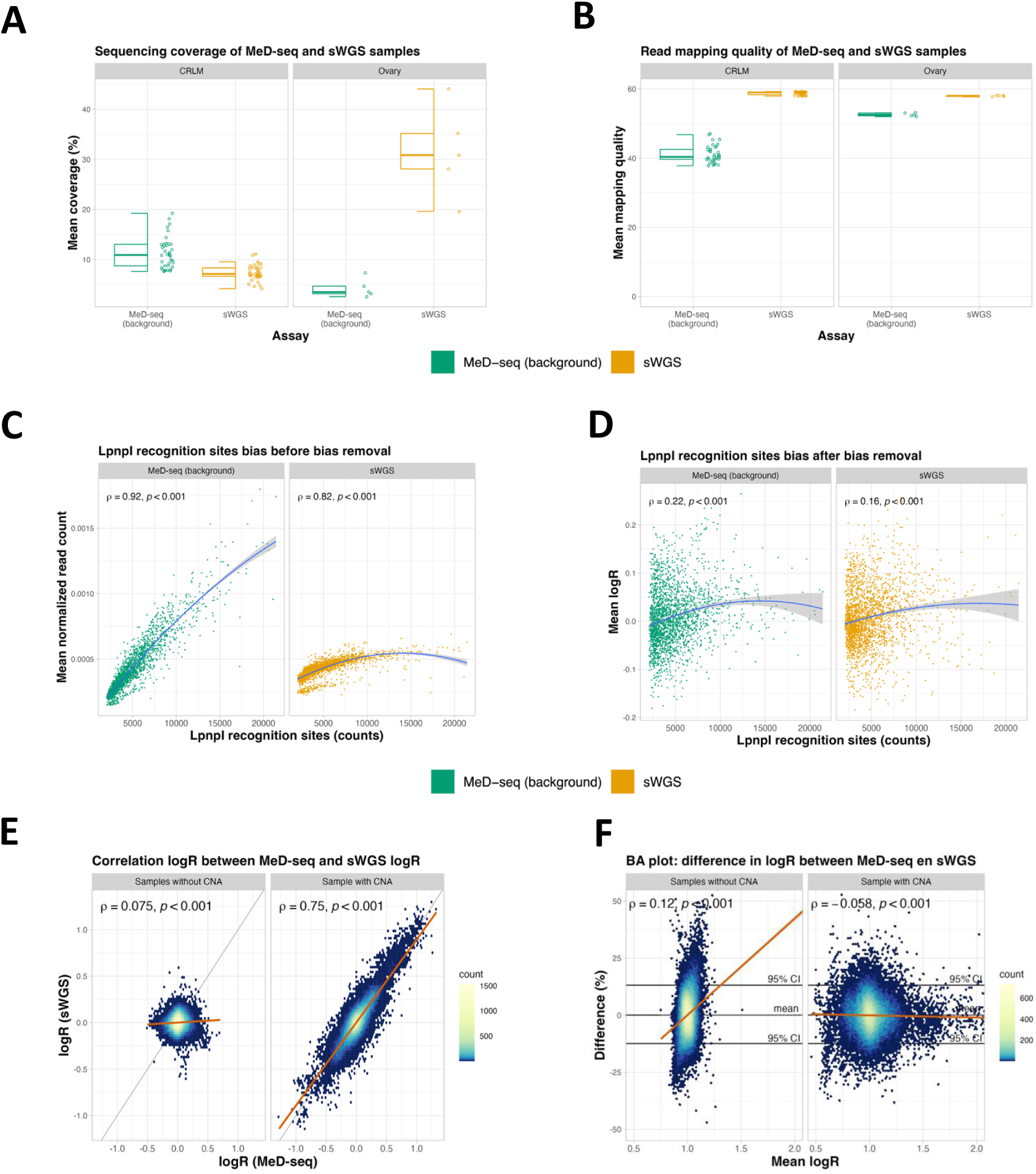
Sequencing statistics, LpnPI bias removal, and strong logR correlation in MeD-Seq data. **A, B)** Scatter and box plots compare sequencing coverage (**A**) and read mapping quality (**B**) per sample (points) for MeD-seq background reads and sWGS data (colors) separately for 38 CRLM and five ovarian cancer samples. **C, D)** Scatter plots show the 1Mb genomic bins (points), normalized read counts (before correction, **C**) and logR (after correction, **D**) versus LpnPI site counts (x-axis) for MeD-seq and sWGS (colors). Blue lines indicate second-order regression, with the shaded area indicating the standard error of the estimate. With Spearman correlation and corresponding p-value in top left. **E)** Heatmaps compare logR values from MeD-seq and corresponding sWGS, split by CNA presence in the sample. With Spearman correlation and corresponding p-value in top left. **F)** Heatmaps show Bland-Altman plots for logR differences, split by CNA presence. With simple linear regression lines, 95% confidence intervals, and Spearman correlation.

We observed a strong correlation between the per bin read counts and both GC content (Spearman ρ = 0.96, p<0.001) and LpnPI recognition site counts (Spearman ρ = 0.92, p<0.001), for MeD-seq background reads, which could negatively impact the identification of CNs (Figure 1C-D, Figure Supplemental Figure 3). Additionally, GC content and LpnPI recognition site counts were highly correlated as well (ρ = 0.95, p < 0.001) (Supplemental Figure 4). To address these biases, we applied LOESS regression, and used a separate panel of MeD-seq normals to generate normalized read count ratios (logR). This approach reduced both GC bias (ρ = 0.11, p<0.001) and LpnPI recognition site bias (ρ = 0.22, p<0.001) in the logR values from MeD-seq (Figure 1D, Supplemental Figure 3).

Next, we compared MeD-seq logR values to sWGS across the same genomic bins and samples. We observed strong correlation (ρ=0.75, p < 0.001) and high agreement (mean relative difference: 0.08%, 95% CI: [-0.04%, 0.19%]), which supports the applicability of MeD-seq logR values for ichorCNA model fitting (Figure 1E-F).

### Copy number profiles and ctDNA load estimates from MeD-seq are concordant with those from sWGS

We fitted ichorCNA using both MeD-seq background reads and paired sWGS data of 43 samples. A representative example output for a cancer sample is shown in Figure 2A,B. We detected a non-zero TF (range: [15%-43%] in the same 21 cancer samples, and we observed a strong correlation between TF-estimates (ρ=0.97, p < 0.001) (Figure 2C). Additionally, we performed BA-analysis which showed high agreement between TF-estimates (mean relative difference: -0.43%, 95%CI: [-1.5%, 0.71%]) (Figure 2C). Using the methylated reads instead of the background reads led to significantly worse CNV profiles and tumor fraction estimates (supplemental figure 5-7), implying that the background reads contain an independent copy-number driven tumor-derived signal.

**Figure 2.**
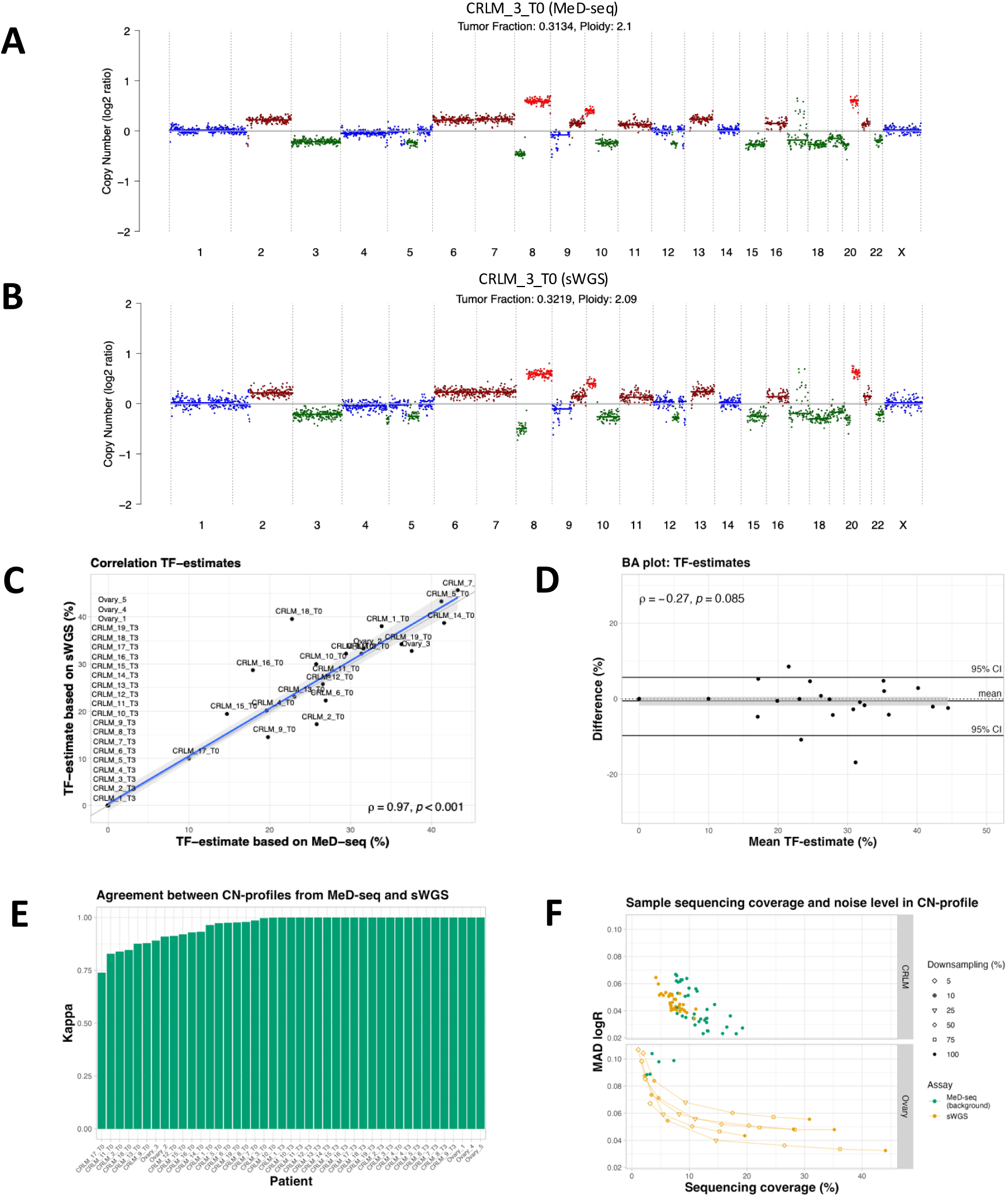
TF Estimates and CN Profiles from MeD-seq Are highly Comparable to sWGS. **A, B)** CN profiles based on MeD-seq (A) and sWGS (B) from patient CRLM_3_T0, showing logR values (y-axis) and predicted CN states (colors) for each genomic bin (x-axis). **C)** Scatter plot of tumor fraction (TF) estimates from MeD-seq (x-axis) and sWGS (y-axis) with Spearman correlation based on non-zero TF samples and corresponding p-value. **D)** Bland-Altman plot showing mean TF estimates (x-axis) versus differences between MeD-seq and sWGS TF estimates (y-axis), with bootstrapped difference (solid lines) and confidence intervals of the mean difference (grey box). **E)** Bar plot of Cohen’s kappa values (y-axis) for CN-profile agreement between MeD-seq and sWGS per patient (38 CRLM, 5 ovary). **F)** Median Absolute Deviation (MAD) in logR values (y-axis) versus sequencing coverage (x-axis) for MeD-seq and sWGS. Down-sampled sWGS data from five high-coverage ovary samples are shown with connected symbols and MAD values.

As an independent validation of the estimated TF, mutation-based variant allele frequencies (VAFs) were available for 19 CRLM samples analysed by targeted colon-specific NGS panel containing 12 genes. A weak to moderate correlation between these mutation-based Variant Allele Frequencies (VAFs) for both MeD-seq (ρ = 0.25, p = 0.3) and sWGS-based TF estimates (ρ = 0.44, p = 0.06) was observed (supplemental figure 8). In two samples, no mutations were detected using the NGS panel, whereas a non-zero TF was detected using ichorCNA both in MeD-seq and sWGS data, indicating these samples contained ctDNA but the somatic mutations present were not covered by the small NGS panel. Copy-number based TF estimates were generally higher than VAFs, likely due to the MeD-seq experimental protocol selecting shorter fragments, and the in silico size-selection for sWGS, both of which enrich for ctDNA fragments (Mouliere et al., 2018).

Next, we assessed the inter-rater agreement of CN-profiles by comparing the integer CN-state assigned to each bin. This analysis showed an average Cohen’s kappa of 0.92 (range: 0.74-1.00) for samples with non-zero TF, and perfect agreement for zero TF samples (Figure 2E).

To assess the effect of read depth on the detection of CNVs, we quantified noise levels in CN-profiles by calculating the Median Absolute Deviation (MAD) of logR values (Supplemental Figure 9). The five ovarian cancer samples showed significantly higher noise in MeD-seq CN-profiles compared to sWGS (p=0.008, Mann-Whitney U test), which we hypothesized was due to higher sequencing coverage in sWGS. To test this, we down-sampled the ovarian sWGS samples to match the MeD-seq coverage, after which similar noise levels were observed (mean MAD, MeD-seq: 0.096, sWGS: 0.092). This provides further evidence that the observed noise differences stem from sequencing coverage and indicates that the quality of MeD-seq CN-profiles is similar to those based on sWGS. (Figure 2F).

Lastly, we compared the distribution of observed CN alterations based on MeD-seq and sWGS in CRLM and ovarian patients (supplemental figure 10, 11). Aggregated CRLM profiles were highly similar between MeD-seq and sWGS data and revealed common copy number gains in chromosomes 7, 8q, 13, and 20, as well as losses in chromosomes 8p and 18, consistent with chromosomal alterations frequently observed in metastatic colorectal cancer (Mendelaar et al., 2021).

### Accounting for local CN-Alterations improves Differential Methylation Modelling

After validating CN profiles and TF estimates from MeD-seq, we analysed the methylation profiles of 27,931 CpG islands in 120 CRLM samples, comparing them to 31 HBDs. We evaluated two models: a CN-naïve Differential Methylation Model (4) and a CN-informed model, which incorporates CN alterations, using the ichorCNA logR values (equation 5), to adjust for changes in expected methylated read counts (Figure 3A, Supplemental figure 12).

**Figure 3.**
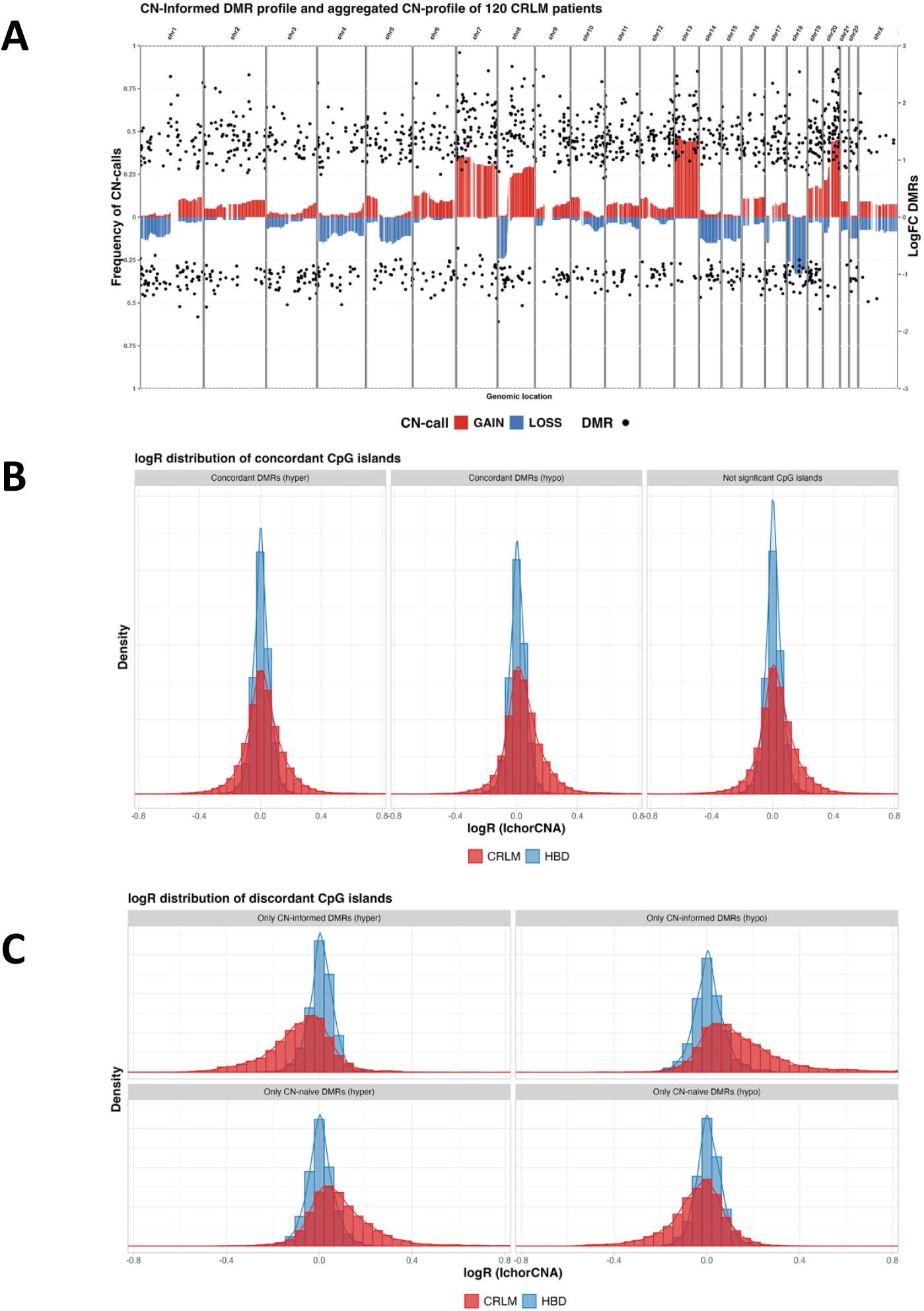
CN-Informed Differential Methylation Analysis Identifies Additional DMRs and Removes CN-Affected CpG islands. **A)** Scatterplots of differentially methylated regions (DMRs) (points), representing significantly differentially methylated CpG islands (DMRs) (FDR < 0.05) identified by the Copy Number (CN) informed Differential Methylation Model. The plot shows log Fold Change (logFC) (right y-axis) across genomic locations (x-axis). Bars indicate aggregated copy number (CN) profiles, with CN gain and loss frequencies in the same dataset (left y-axis) represented by color. **B, C)** Histograms and density plots showing logR value (x-axis) distributions for each CpG islands in CRLM patients and Healthy Blood Donors (HBDs) (colors) indicating the effect of CN alterations on DMRs. **B)** CpG islands classified as hypermethylated, hypomethylated, or not significant by both CN-naïve and CN-informed models. **C)** CpG islands uniquely identified as DMRs by either the CN-informed or CN-naïve model, separated by hypermethylated and hypomethylated DMRs.

The CN-informed model identified 1,482 differentially methylated regions (DMRs) (FDR-adjusted p-value<0.05), including 1,029 (69%) hypermethylated and 453 (31%) hypomethylated regions (Figure 3A-B). The CN-naive model identified 1,589 DMRs, including 1,132 (71%) hypermethylated and 457 (29%) hypomethylated regions. Hypermethylated DMRs were primarily found in CN-gained regions on chromosomes 7, 8q, 13, and 20, while hypomethylated DMRs were enriched in CN-losses, particularly on chromosomes 8p and 18 (Supplemental Figure 13). The lower deviance statistic of CpG islands in the CN-informed model compared to the CN-naïve model suggests an improved model fit (Wilcoxon Signed-Rank Test, p<0.001) (Supplemental Figure 14). Among the DMRs identified, 1,370 were concordant between both models, with a strong correlation in logFC values (r=0.99, p<0.001) (Supplemental Figure 15, table 1). However, the CN-informed model detected 112 additional DMRs and excluded 219 that were likely driven by CN alterations.

**Table 1.**
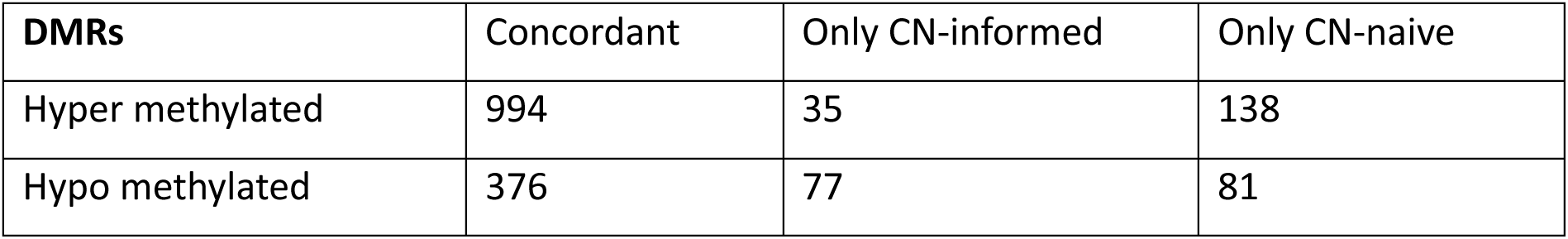
Number of DRMs detected by CN-informed and CN-naïve differential methylation analysis.

| DMRs | Concordant | Only CN-informed | Only CN-naïve |
| --- | --- | --- | --- |
| Hyper methylated | 994 | 35 | 138 |
| Hypo methylated | 376 | 77 | 81 |

DMRs not affected by CN alterations were consistently identified by both the CN-naïve and CN-informed models and exhibited similar logR distributions in healthy individuals and CRLM patients (Figure 3B). In contrast, DMRs influenced by CN alterations showed altered logFC values due to these genomic changes. As a result, the CN-informed model was able to detect additional DMRs that were obscured in the CN-naïve analysis. Furthermore, the CN-informed model removed false-positive DMRs identified by the CN-naïve model by adjusting for CN variation using the sample-specific logFC values of each CpG island (Figure 3C).

### CNV analysis identifies ctDNA-high samples leading to increased power for differential methylation

To investigate the impact of tumor fraction (TF) on methylation profiles, we categorized the 120 CRLM samples into 65 high-TF (≥5%) and 55 low-TF (<5%) samples based on IchorCNA estimates. The CN-informed model was applied to each subset, using the 31 HBDs as controls to identify DMRs. Analysis of the 55 low-TF samples yielded only a single hypermethylated DMR, indicating limited power to detect differential methylation in this group. To determine whether this was due to sample size, we performed bootstrapping by randomly selecting 55 samples from the full cohort to match the size of the low-TF subset. This analysis resulted in the detection of 44 DMRs, suggesting that tumor load and thus the effect size rather than sample size is the limiting factor. Moreover, bootstrapping 55 samples from the high-TF subset produced a much stronger signal, detecting a total of 1,016 DMRs—including the previously identified regions—and highlighting the increased statistical power when analyzing only high-TF samples (Table 2, Figure 4A).

**Figure 4.**
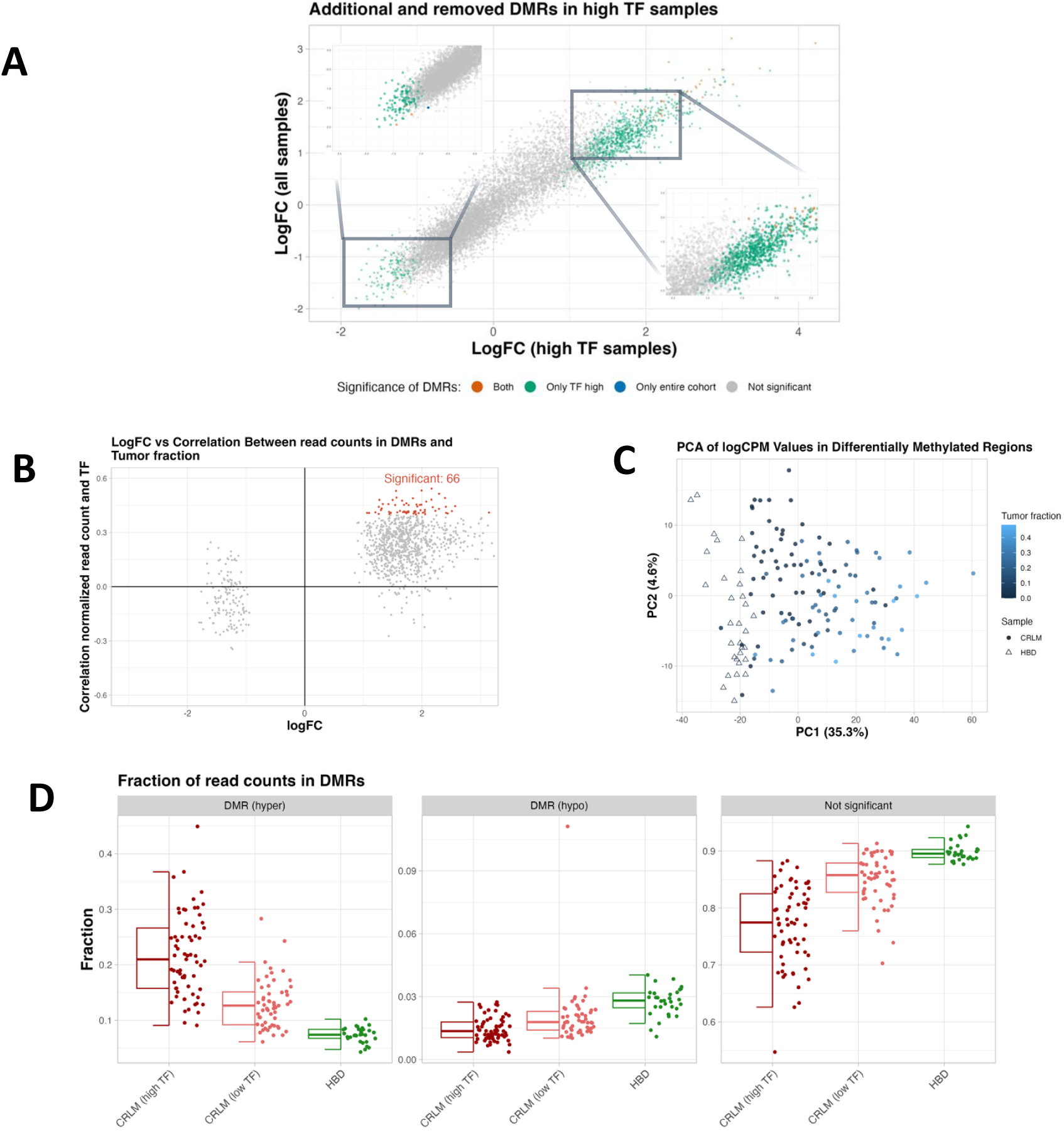
Differential Methylation Analysis is Influenced by TF in samples. **A)** Scatterplot comparing log Fold Change (log FC) values from differential methylation analysis based on high Tumor Fraction (TF) samples (x-axis) and entire cohort (y-axis). Colors indicate whether CpG islands were classified as Differentially Methylated Regions (DMRs) based on both datasets, only one, or neither. **B)** Scatterplot showing the Spearman correlation (y-axis) between normalized read counts and tumor fraction (TF) across high-TF CRLM samples for DMR (points), plotted against the corresponding log fold-change (logFC) (x-axis). Points are colored red if the correlation with TF is statistically significant (FDR-adjusted p < 0.05). **C)** Principal Component Analysis (PCA) of all cohort samples based on normalized read counts in DMRs detected using only the high-TF samples. Each point represents a sample, colored by its ichorCNA-estimated tumor fraction (TF). Point shapes distinguish Healthy Blood Donors (HBDs) from CRLM patients. **D)** Scatter and box plots showing the fraction of total reads (y-axis) per sample (points), grouped by sample category on the x-axis: Healthy Blood Donors (HBDs), CRLM patients with low tumor fraction (TF < 5%), and high tumor fraction (TF ≥ 5%). Separate panels display read fractions in hypermethylated DMRs (left), hypomethylated DMRs (middle), and all not significantly differentially methylated CpG islands (right). Colors correspond to sample groups.

**Table 2.** Number of DRMs detected entire cohort, high TF samples, low TF samples when bootstrapping to 55 samples.

| DMRs | Entire cohort | High TF samples | Low TF samples |
| --- | --- | --- | --- |
| Hyper methylated | 40 | 877 | 0 |
| Hypo methylated | 4 | 139 | 1 |

Since DMR calling appeared to be influenced by the tumor fraction (TF) present in a sample, we proceeded by focusing on the DMRs identified using the high-TF subset. For each DMR, we determined the Spearman correlation between the normalized, CN-corrected, and log-transformed read counts and the corresponding ichorCNA TF estimates. A one-sided test was applied, using the sign of the logFC to determine the direction of the expected correlation. This identified 66 significant positive correlations (FDR-adjusted p < 0.05), between tumor fraction and normalized reads counts in DMRs (Figure 4B, Supplemental Figure 16). Additionally, principal component analysis (PCA) of normalized methylation counts across all DMRs revealed clear separation between CRLM and HBD samples. Notably, the first principal component (PC1) showed a postive correlation with tumor fraction (ρ = 0.64, p < 0.001), indicating a gradient driven by tumor-derived methylation signals (Figure 4C).

To explore the distribution of tumor-associated methylation changes, the fraction of total reads mapping to hyper- or hypomethylated DMRs was calculated for each sample. As expected, the high-TF samples had a greater proportion of reads in hypermethylated DMRs (p<0.001) and a reduced proportion in hypomethylated DMRs (p<0.001) compared to HBDs. Interestingly, although the DMRs were identified using only the high-TF samples, the low-TF samples also showed a similar trend—with elevated read fractions in hypermethylated DMRs (p<0.001) and decreased fractions in hypomethylated DMRs (p<0.001) relative to HBDs (Figure 4D). Lastly, we examined the relationship between tumor fraction estimates and the fraction of reads mapping to hypermethylated DMRs, revealing a significant positive correlation (ρ = 0.40, p = 0.001) and a moderate negative correlation (ρ = -0.24, p = 0.52) for the hypomethylated DMRs. Together, these findings demonstrate how our multi-modal analysis of MeD-seq data enables more accurate detection of tumor-associated methylation changes, leveraging samples with a high circulating tumor fraction to improve detection even in samples with low tumor burden.

## Discussion

In this study we evaluated a novel approach for simultaneous methylation- and chromosomal CN profiling, and TF estimation from cfDNA using data generated by our previously developed MeD-seq assay (Boers et al., 2018; Deger et al., 2021). We showed that it is feasible to extract CN-profiles and TF-estimates from MeD-seq that are highly comparable to those obtained from sWGS. Moreover, we combined CN and TF information with methylation profiling, which enables the identification of additional DMRs related to tumor load in CRLM patients.

We leveraged background reads of MeD-seq data to derive CN profiles and TF estimates, using the ichorCNA model, without altering the experimental protocol of the assay. This reduces the need for additional biological material and adds relevant information to readily performed MeD-seq experiments. Even though MeD-seq data contained a bias due to LpnpI digestion, this bias can consistently be reduced resulting in CN-profiles that can be constructed using a sample coverage of 2.5% from a total of 30 million reads sequenced per sample. The resulting CN-profiles provide per-sample genomic insights that could support further characterization of CRLM patients (Eikenboom et al., 2023; Mendelaar et al., 2021; Tan et al., 2022; van Rees et al., 2023).

An additional use of CN-profiles data is its integration into the Differential Methylation Model. In MeD-seq, differential methylation is based on counts of LpnPI-digested DNA fragments, which can be influenced by chromosomal alterations, such as gains and losses. Since CN alterations and varying TFs are common in cancer, incorporating this information helps reduce the potential for misinterpreting differential methylation signals due to underlying CN changes (Adalsteinsson et al., 2017; Martínez-Jiménez et al., 2022). To address this, we included read count ratio values as normalization in the modeling process, which resulted in the detection of additional DMRs and the removal of likely false DMRs caused by CN alterations.

The amount of ctDNA can vary considerably among cancer patients, even within the same disease stage (Cristiano et al., 2019). By jointly profiling CNVs, TF estimation, and methylation, we gain an independent strategy for selecting samples from patients with sufficient ctDNA for DMR analysis within the same experiment. Additionally, CN-based ctDNA load estimates can be correlated to the methylation profile of DMRs to assess whether the changes methylation are indeed tumor-related. We found that focusing on ctDNA-high samples resulted in a larger number of DMRs significantly associated with tumor load. Additionally, using the DMRs detected in high-TF samples, we were able to distinguish low-TF samples from HBDs, suggesting that MeD-seq-derived DMRs could serve as potential ctDNA biomarkers. Taken together, these results show how our new multi-modal analysis of MeD-seq data can lead to improved detection of tumor-derived methylation changes.

The ability to simultaneously extract methylation and copy number information from a single cfDNA assay like MeD-seq holds promising potential for minimally invasive tumor profiling as it reduces both analysis costs and the amount of cfDNA needed (Kim et al., 2023; Li & Sun, 2024). Our approach may contribute to more refined patient stratification and biomarker discovery, as stratification based on tumor fraction, can help identify patients with sufficient ctDNA burden for reliable epigenetic analysis—especially relevant in early detection or monitoring scenarios where ctDNA levels are generally low. Moreover, incorporating CN-informed modeling reduces false positives by correcting for genomic alterations that might confound methylation signals, thereby improving specificity of detected DMRsThe methodological advances presented here represent a step toward more informative, interpretable, and affordable cfDNA-based diagnostics. Expanding MeD-seq applications across diverse cancer types will be essential to further evaluate its clinical utility and potential as a cost-effective, non-invasive diagnostic tool.

## Abbreviations

CRLM: ColoRectal cancer with Liver Metastasis
MeD-seq: Methylated DNA Sequencing
sWGS: shallow Whole Genome Sequencing
CN: Copy Number
TF: Tumor Fraction
logR: Log2 of the ratio of reads counts (ichorCNA)
logFC: log Fold Change of read counts (edgeR)
VAF: Variant Allele Frequency
DMR: Differentially Methylated Region (CpG island with p-value < 0.05)
MAD: Median Absolute Deviation

## Acknowledgements

We would like to thank Dr. Ingrid Boere and Dr. Heleen van Beekhuizen for their valuable support in collecting and providing the ovarian cancer patient samples used in this study.

