## Supplemental figures for "Improved circulating tumor DNA profiling by simultaneous extraction of DNA methylation and copy number information from Methylated DNA Sequencing data (MeD-seq)"

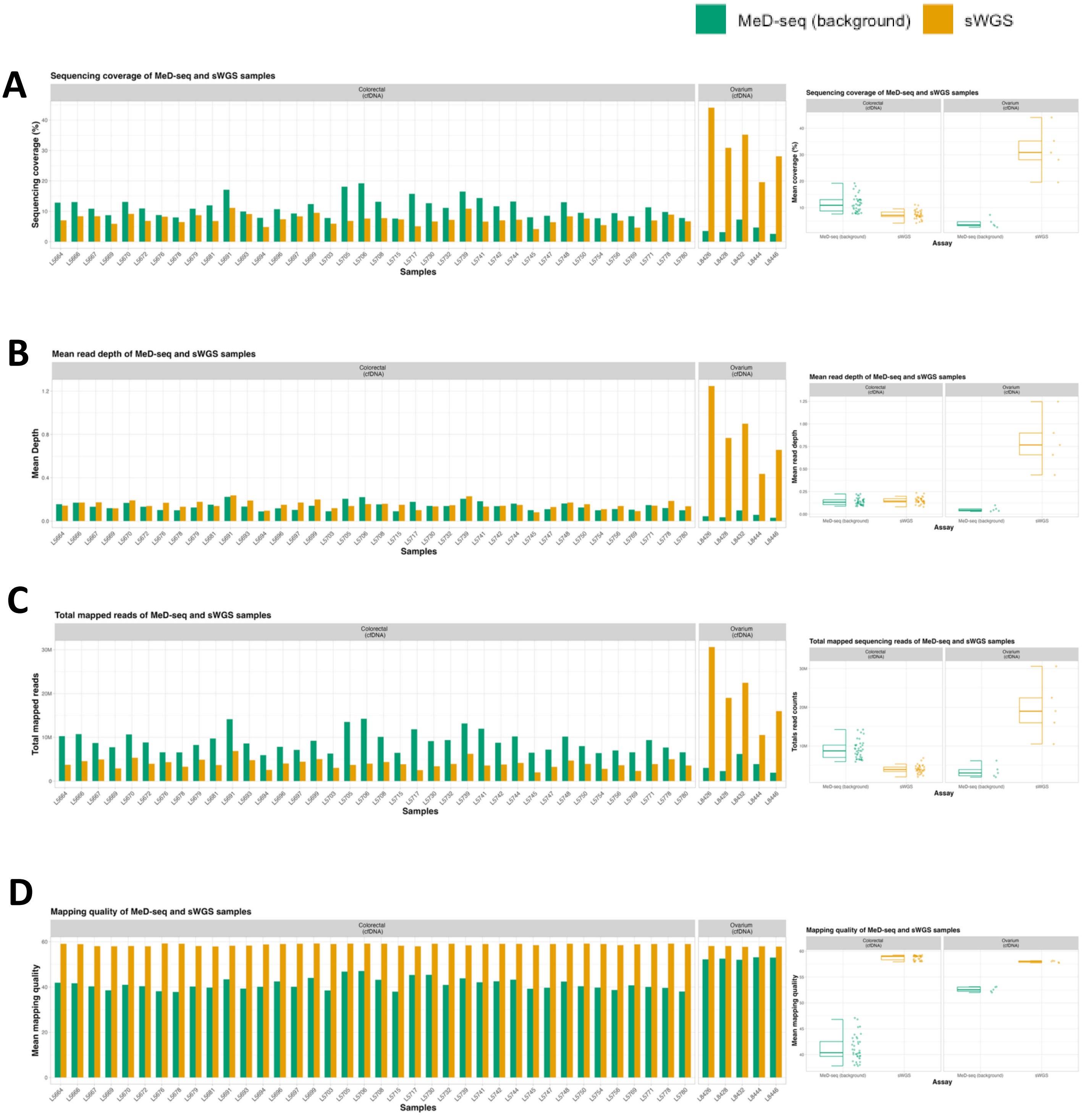


**Supplemental figure 1: Sequencing statistics of MeD-seq and sWGS samples**
Bar plots (per sample) and box plots (per cancer type) depicting sequencing coverage (A), mean read depth (B), total mapped reads (C), and mapping quality (D), shown separately for MeD-seq background reads and sWGS data from the same sample (colors).

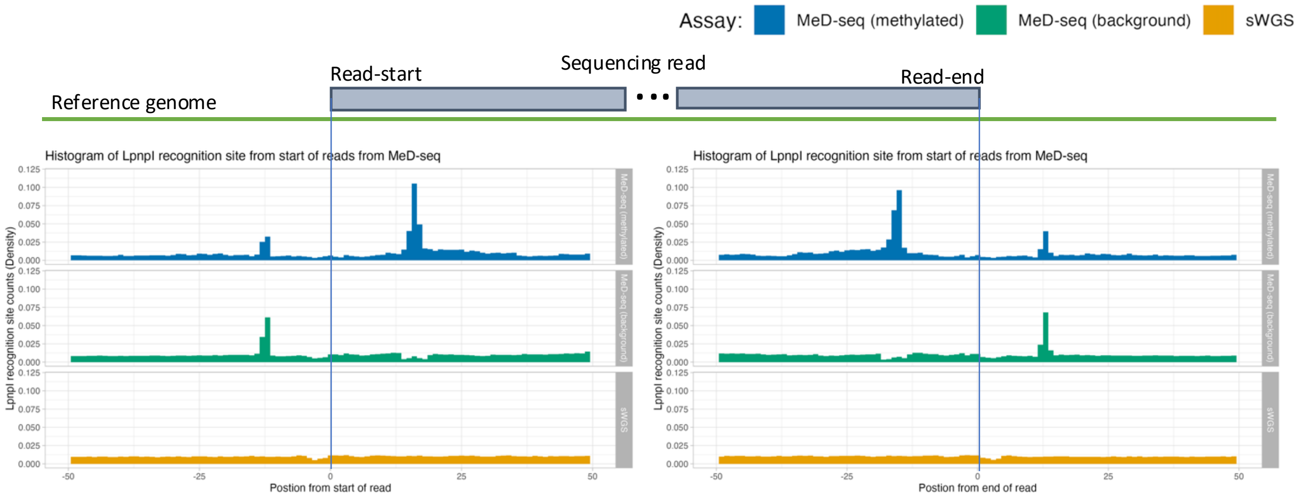


**Supplemental figure 2: LpnpI recognition sites near start and end of sequencing reads.** Histogram showing the occurrence of LpnpI recognition sites (y-axis) near the start (left) and end (right) of 10.000 randomly sampled reads from MeD-seq data and sWGS data (colors) indicating methylated reads (top), background reads (middle) and sWGS reads (bottom) of 5 ovary cancer samples.

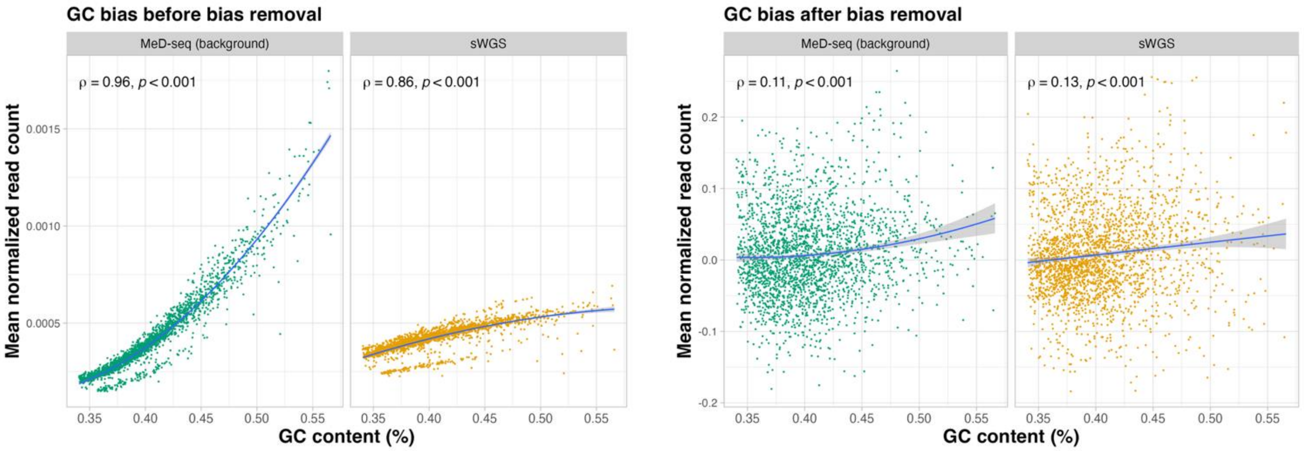


**Supplemental figure 3, removal of GC bias**

Dot plots indicating for each genomic bin (points) the mean normalized read counts before bias removal (y-axis, left) and mean logR after bias correction (y-axis, right), and GC-content (x-axis). Seperatly for MeD-seq and sWGS (colors). The blue line represents second order linear regression line, with the shaded area indicating the standard error of the estimate. The Spearman correlation coefficient and corresponding p-value is depicted in the top left of each plot.

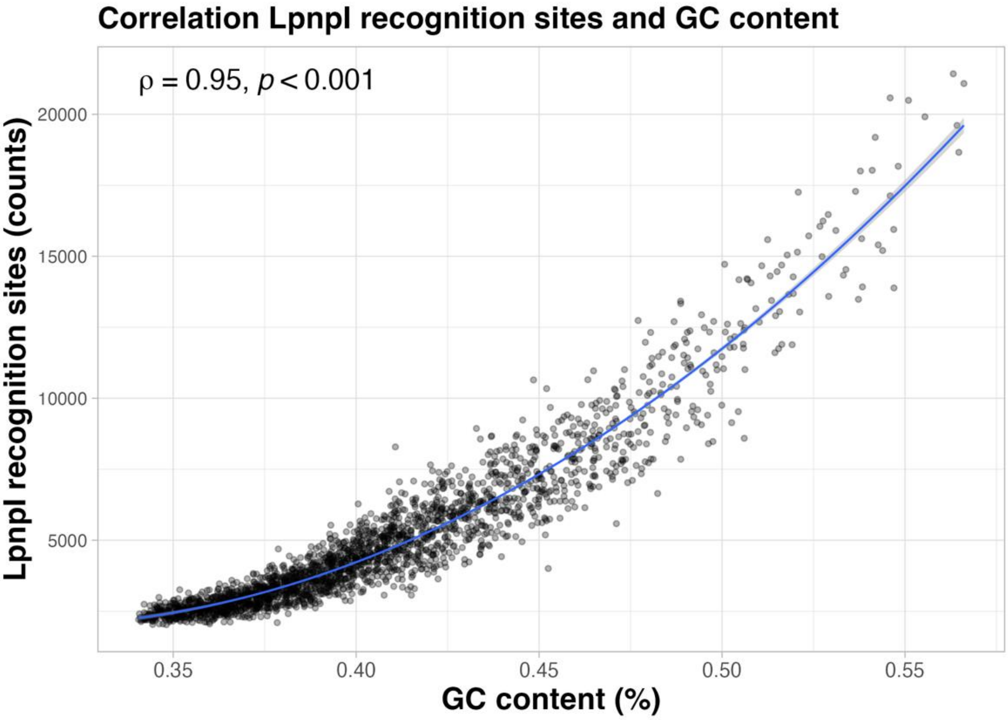


**Supplemental figure 4, correlation between LpnPI recognition sites and GC content**
Scatter plot indicating for each genomic bin (points) the number of LpnPI recognition sites (y-axis) and GC-content (x-axis). The blue line represents second order linear regression line, with the shaded area indicating the standard error of the estimate. The Spearman correlation coefficient and corresponding p-value is depicted in the top left of the plot.

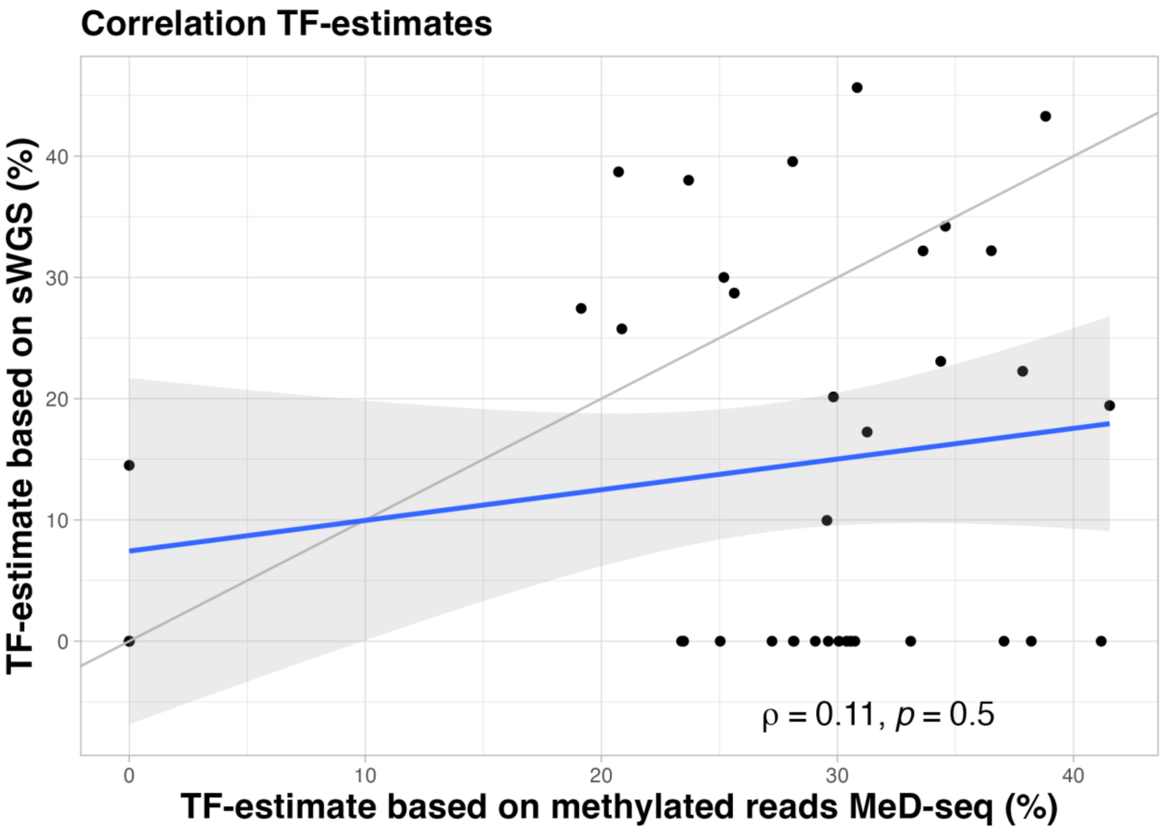


**Supplemental figure 5: Correlation between Tumor Fraction (TF) estimates from MeD-seq methylated reads and sWGS.** Scatterplot of tumor fraction (TF) estimates from methylated reads from MeD-seq (x-axis) and sWGS (y-axis) with Spearman correlation and corresponding p-value.

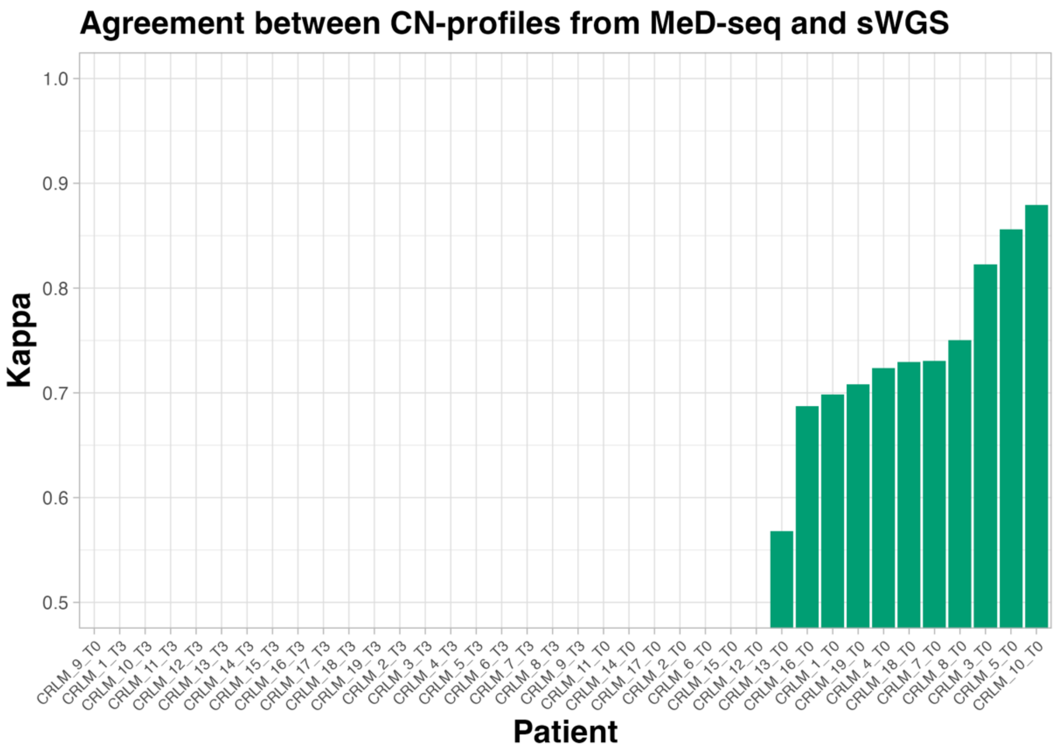


**Supplemental figure 6: Agreement between CN-profiles based on Methylated Reads MeD-seq and sWGS.** Bar plot of Cohen’s kappa values (y-axis) for CN-profile agreement between MeD-seq and sWGS per patient (38 CRLM patients).

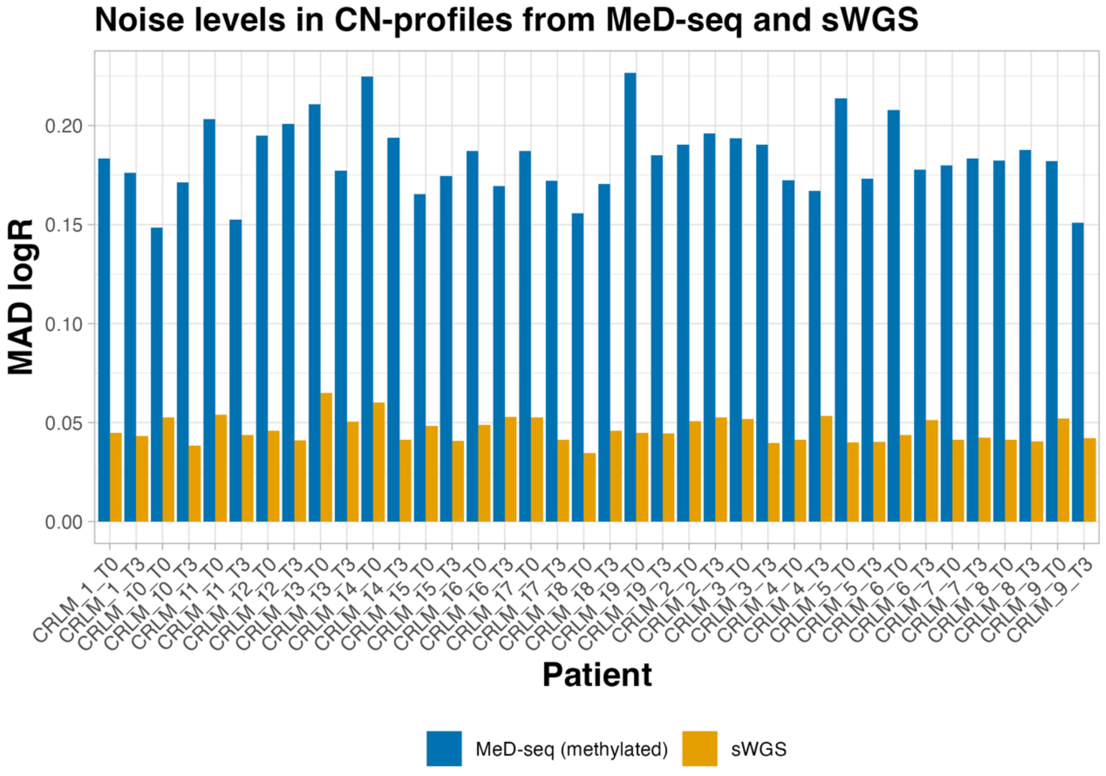


**Supplemental figure 7: Noise levels in CN-profiles based on methylated reads from MeD-seq and sWGS.** Median Absolute Deviation (MAD) in logR values (y-axis) were used to quantify noise in CN-profiles based on MeD-seq and sWGS (colors) in the same patient (x-axis).

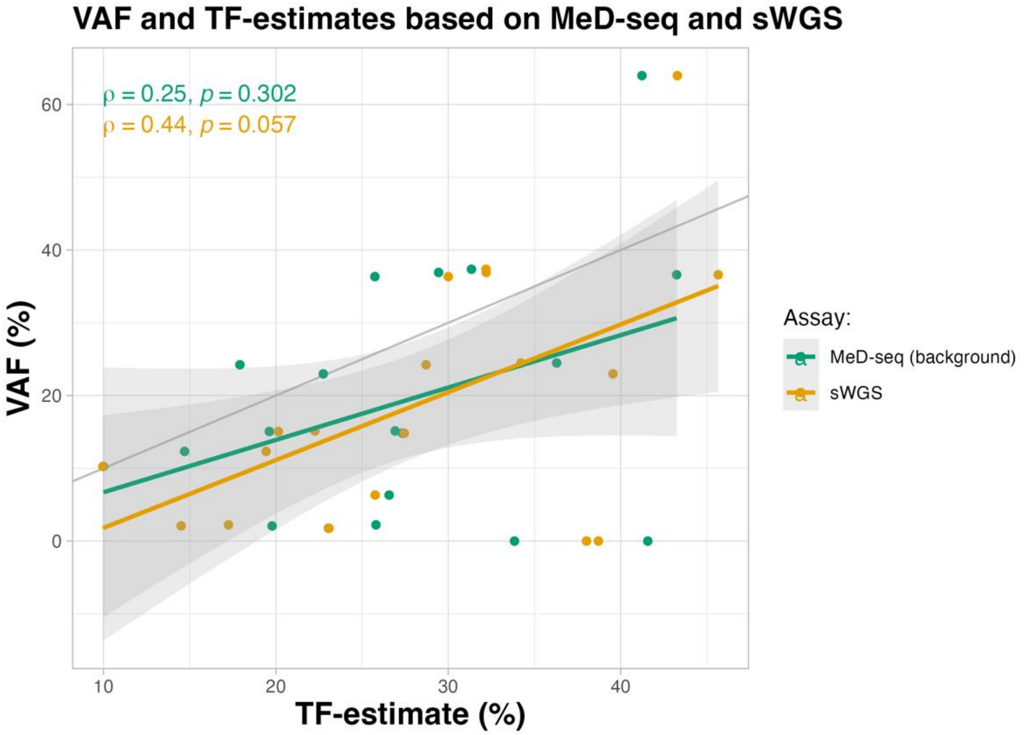


**Supplemental figure 8: Correlation between Tumor Fraction (TF) estimates from MeD-seq, sWGS, and Variant Allele Frequency (VAF).** Scatterplot showing for each sample (points) the Variant Allele Frequency (VAF) (y-axis) and TF estimates from IchorCNA (x-axis) using MeD-seq or sWGS (colors). The solid-coloured lines represents simple linear regression lines, with the shaded area indicating the standard error of the estimate. The Spearman correlation coefficient and corresponding p-value is depicted in the top left .

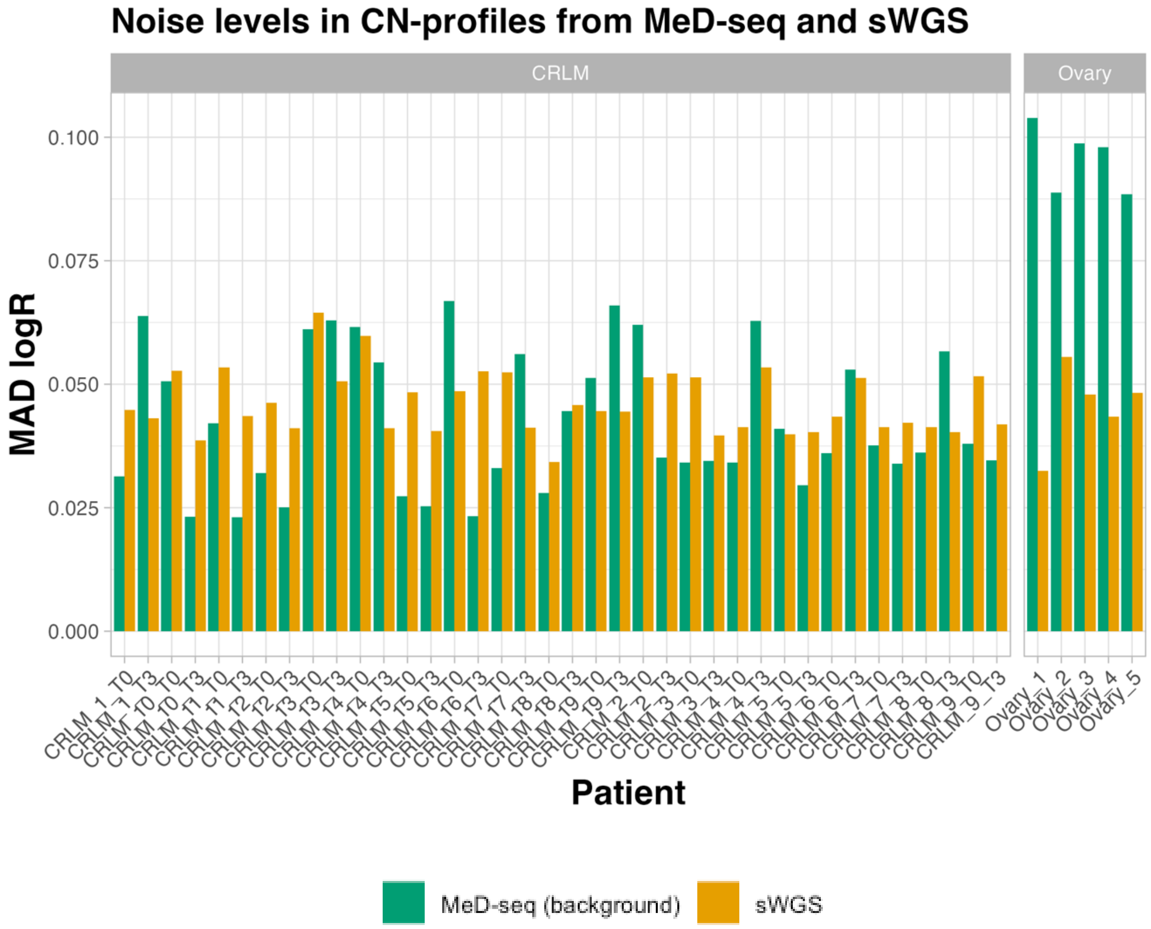


**Supplemental figure 9: Noise levels in CN-profiles based on MeD-seq and sWGS.**
Median Absolute Deviation (MAD) in logR values (y-axis) were used to quantify noise in CN-profiles based on MeD-seq and sWGS (colors) in the same patient (x-axis).

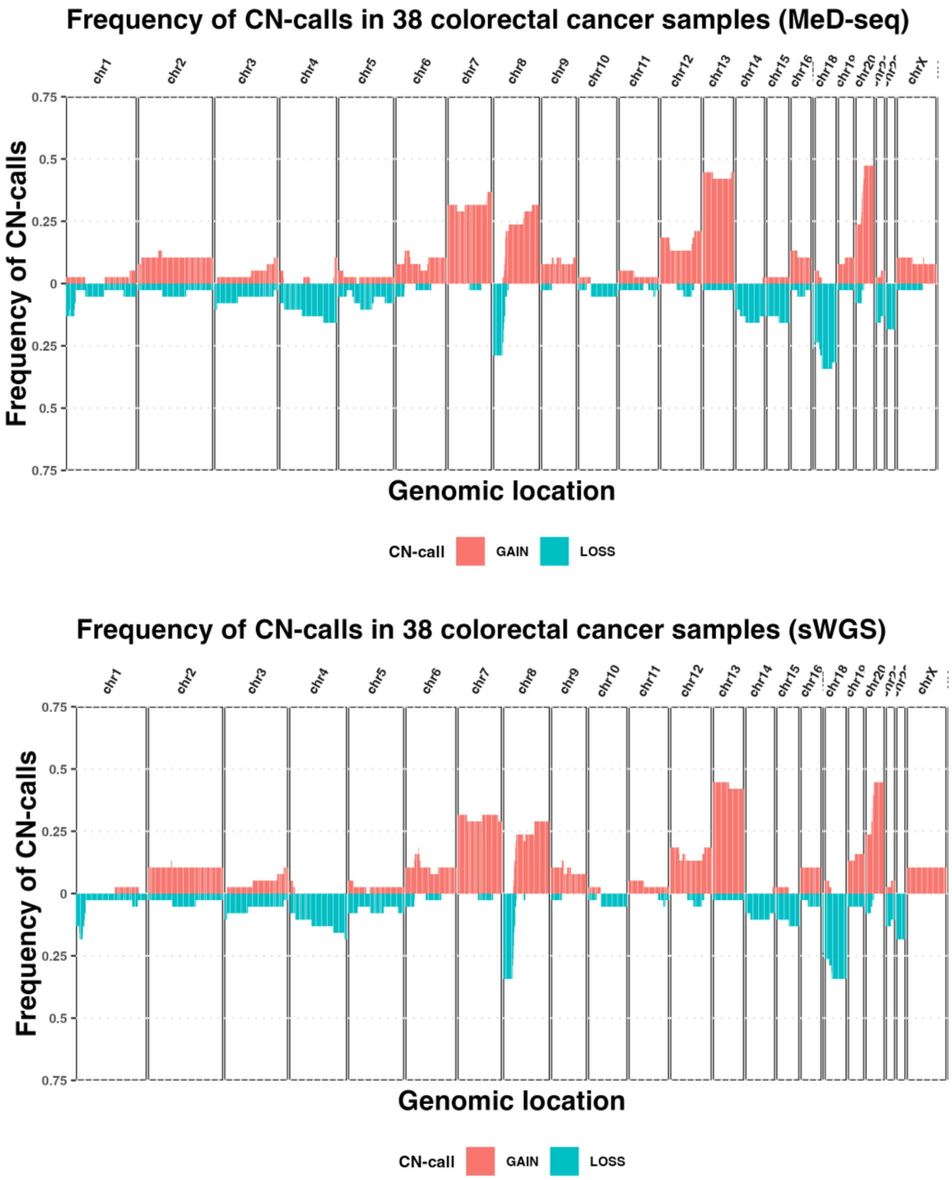


**Supplemental figure 10, Frequency of Gains and Losses in CN-profile based on MeD-seq and sWGS.** Bar plots displaying the frequency (y-axis) of copy number gains and losses (colors) across genomic bins (x-axis) in 38 colorectal cancer patients. The top panel represents aggregated CN-profiles derived from MeD-seq data, while the bottom panel shows aggregated CN-profiles obtained from sWGS.

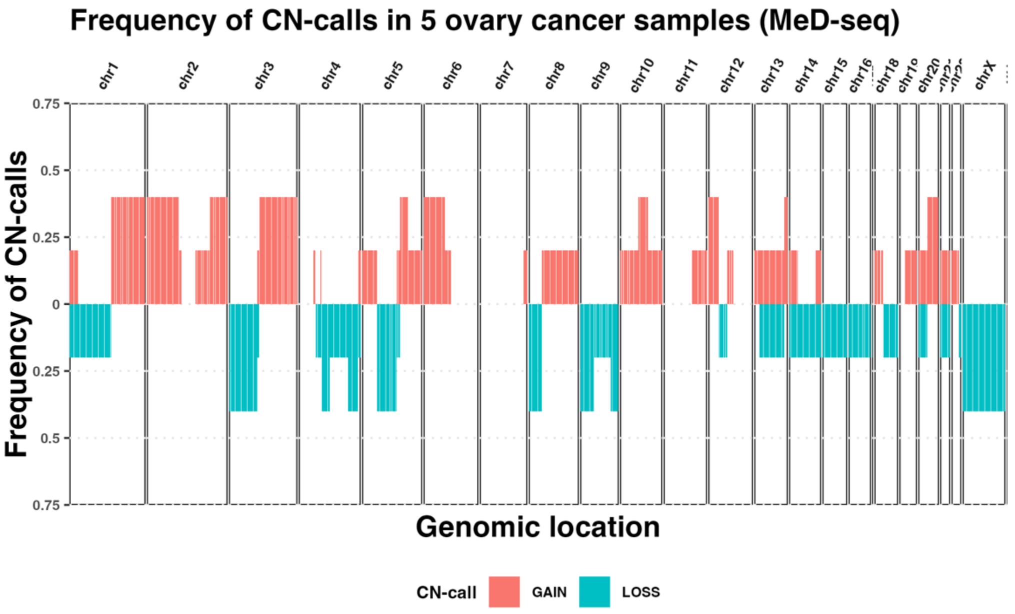


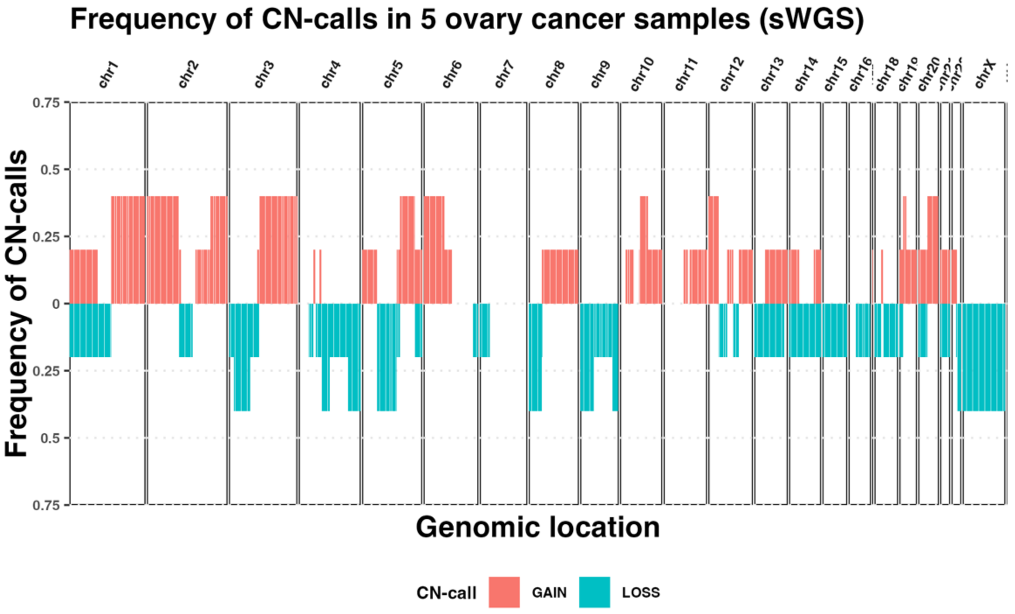


**Supplemental figure 11, Frequency of Gains and Losses in CN-profile based on MeD-seq and sWGS.** Bar plots displaying the frequency (y-axis) of copy number gains and losses (colors) across genomic bins (x-axis) in 5 ovary cancer patients. The top panel represents aggregated CN-profiles derived from MeD-seq data, while the bottom panel shows aggregated CN-profiles obtained from sWGS.

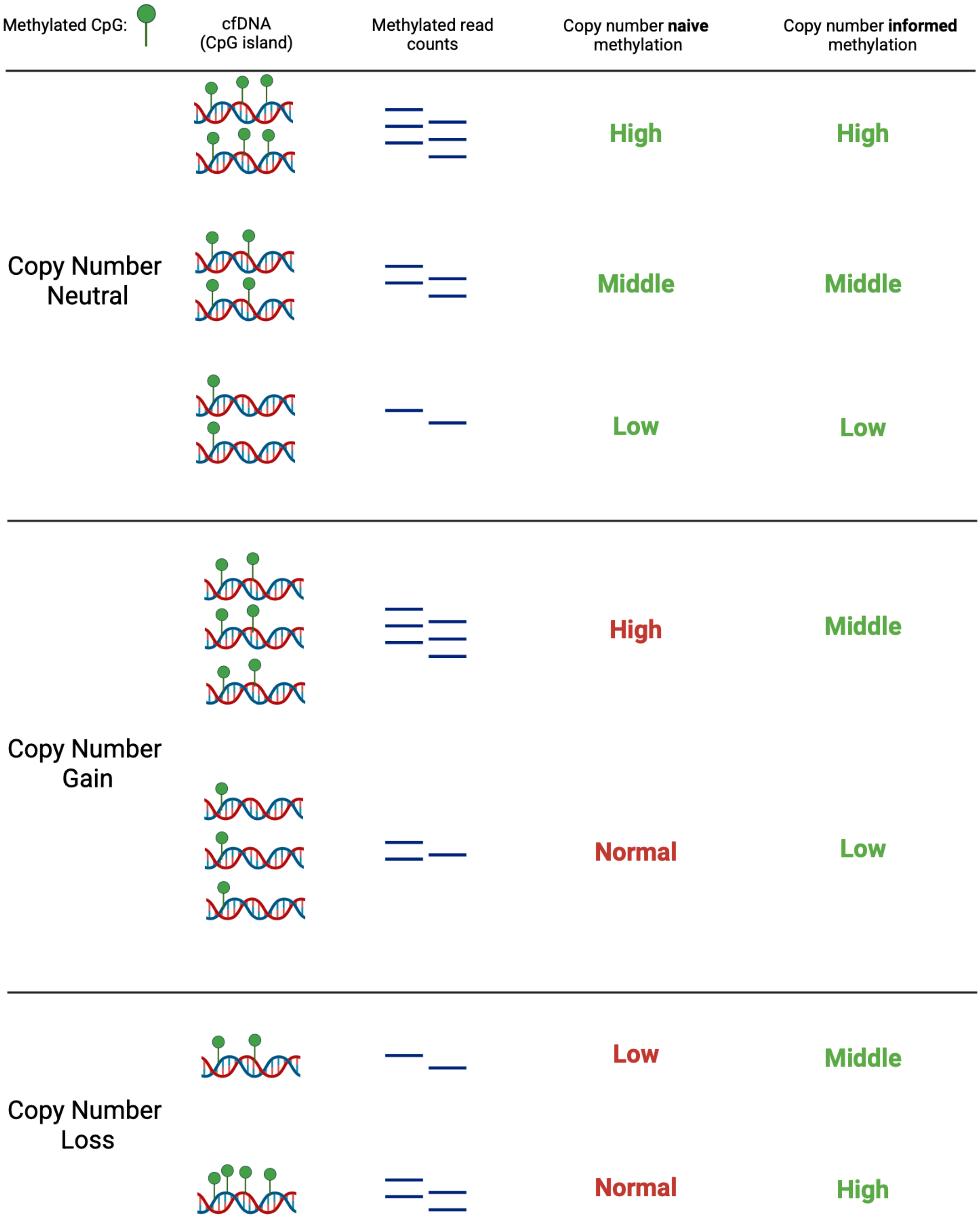


**Supplemental Figure 12: Graphical abstract illustrating the impact of copy number alterations on methylation detection by MeD-seq.**
To account for aberrant read counts in methylation signals caused by copy number gains or losses, copy number information was incorporated into the Copy Number-Informed Differential Methylation Model (CN-informed DMM).

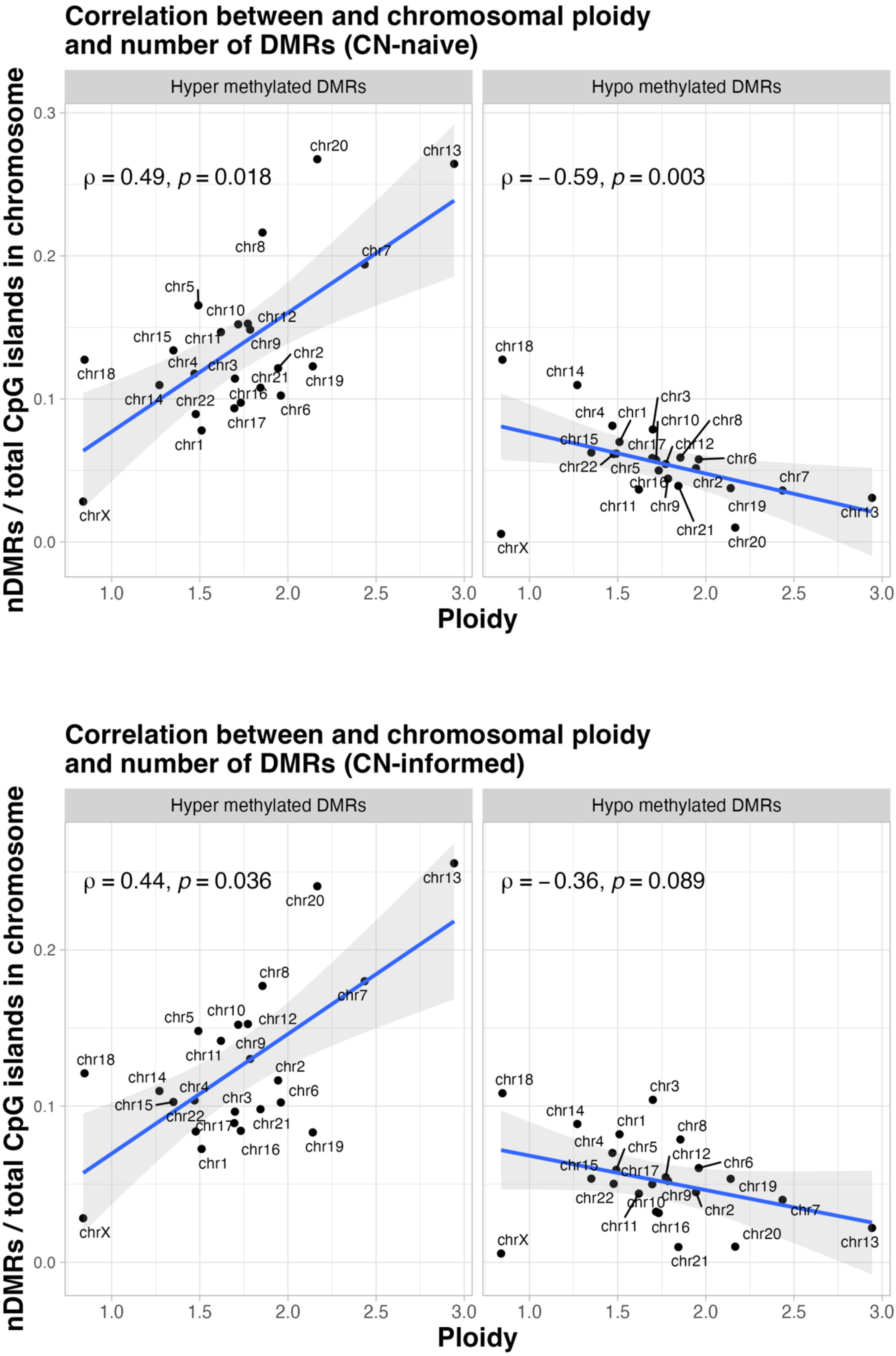


**Supplemental Figure 13: Correlation Between Ploidy and DMR Count**

Scatterplots depicting for each chromosome (points) the median ploidy (x-axis) across 120 CRLM patients and the fraction of CpG islands classified as differentially methylated (y-axis) according to the CN-naïve (top) and CN-informed DMM (bottom).

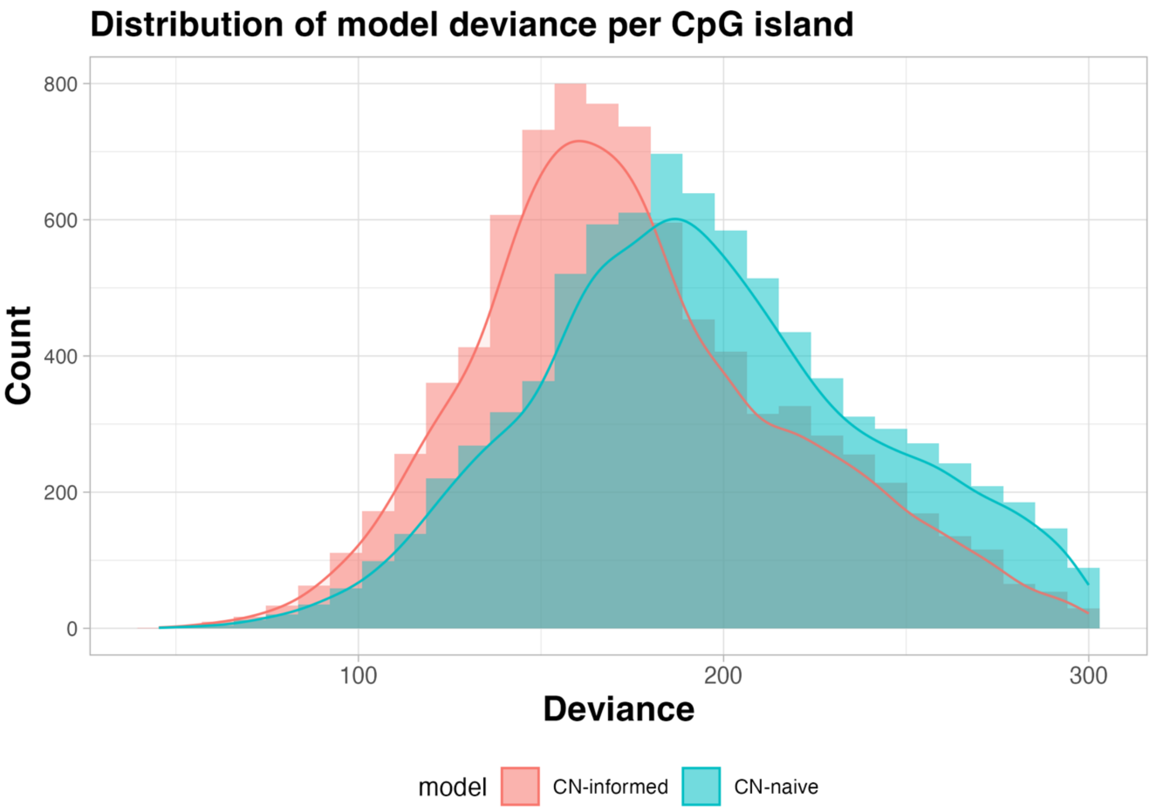


**Supplemental Figure 14**: **Distribution of the deviance statistic for CN-informed and CN-naïve models.** Histograms and densities display the distribution of the deviance statistic (x-axis) for each CpG island, separately for the CN-informed and CN-naïve models (colored).

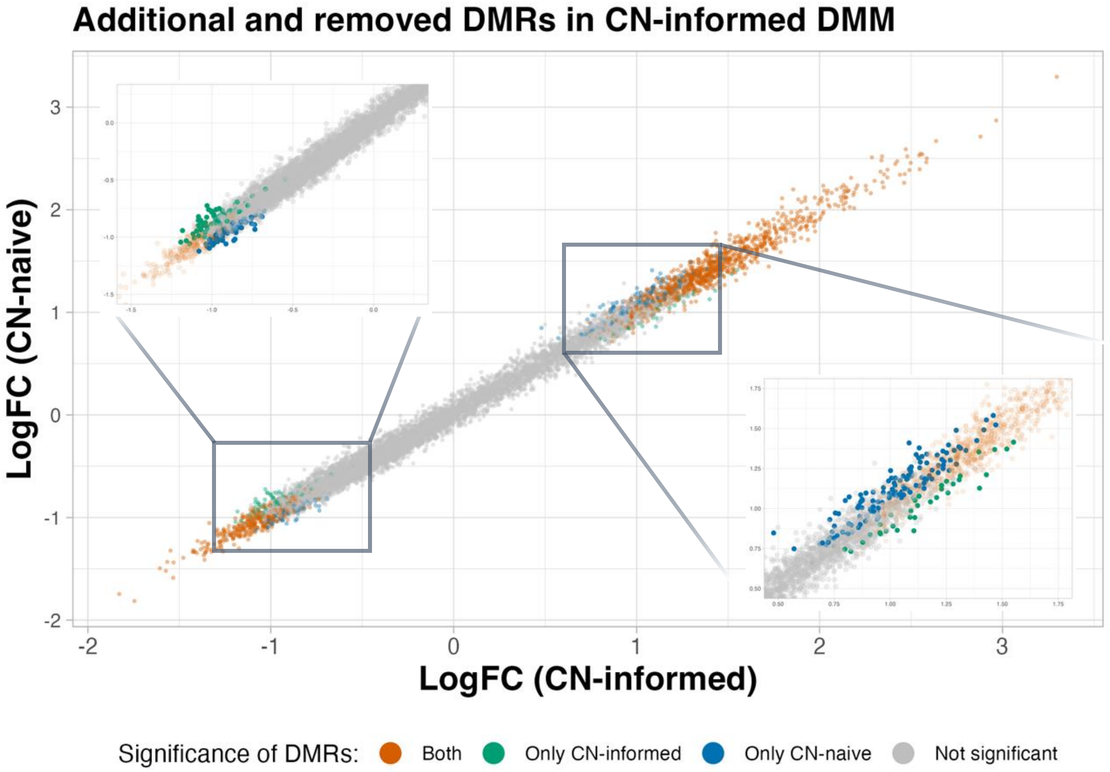


**Supplemental Figure 15**: Scatterplot comparing logFC values from the CN-informed (x-axis) and CN-naïve (y-axis) models. Colors indicate whether CpG islands were classified as DMRs by both models, only one model, or neither.

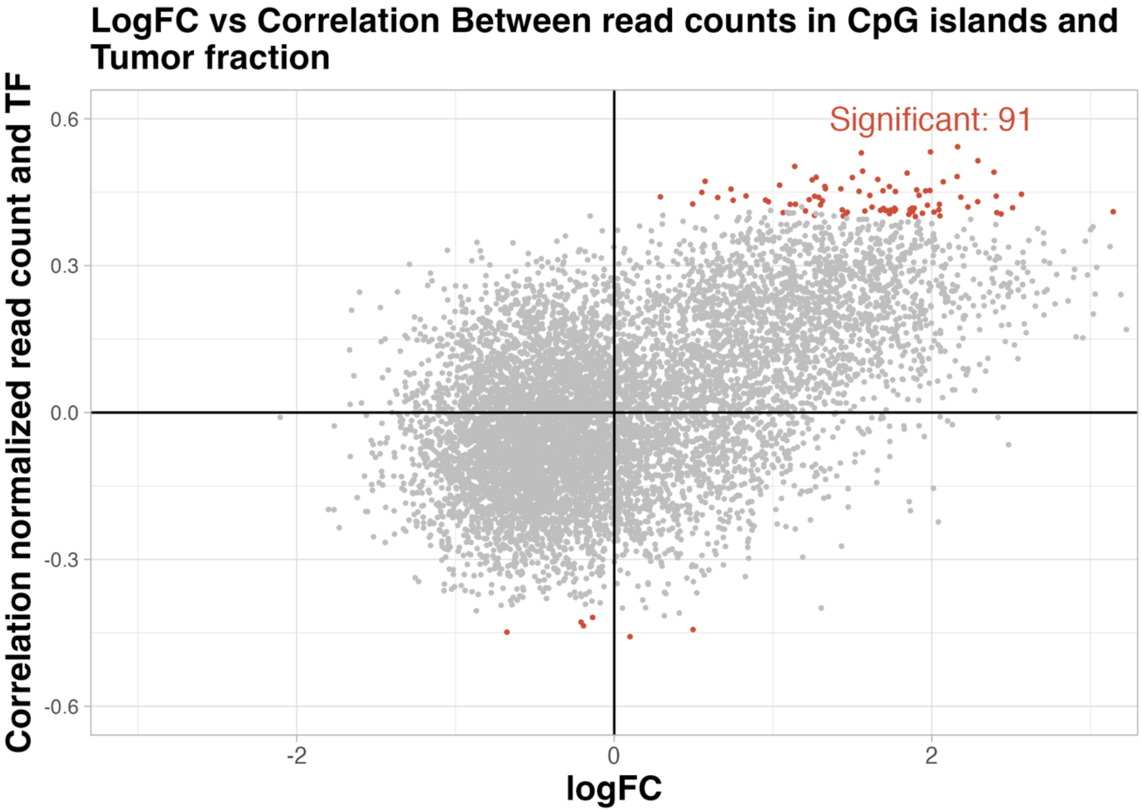


**Supplemental Figure 16: Correlation of CpG Island Read Counts with Tumor Fraction and logFC**
Scatterplot showing the Spearman correlation (y-axis) between normalized read counts and tumor fraction (TF) across high-TF CRLM samples for each CpG island (points), plotted against the corresponding log fold-change (logFC) from the CN-informed DMM (x-axis). Points are colored red if the correlation with TF is statistically significant (FDR-adjusted $p < 0.05$).
